# ZNFX1 is a Novel Master Regulator in Epigenetically-induced Pathogen Mimicry and Inflammasome Signaling in Cancer

**DOI:** 10.1101/2024.10.18.618659

**Authors:** Lora Stojanovic, Rachel Abbotts, Kaushlendra Tripathi, Collin M. Coon, Saranya Rajendran, Elnaz Abbasi Farid, Galen Hostetter, Joseph W. Guarnieri, Douglas C. Wallace, Sheng Liu, Jun Wan, Gennaro Calendo, Rebecca Marker, Zahra Gohari, Mohammed M. A. Inayatullah, Vijay K. Tiwari, Tanjina Kader, Sandro Santagata, Ronny Drapkin, Stefan Kommoss, Jacobus Pfisterer, Gottfried E Konecny, Ryan Coopergard, Jean-Pierre Issa, Boris J. N. Winterhoff, Michael J. Topper, George E. Sandusky, Kathy D. Miller, Stephen B. Baylin, Kenneth P. Nephew, Feyruz V. Rassool

## Abstract

DNA methyltransferase and poly(ADP-ribose) polymerase inhibitors (DNMTis, PARPis) induce a stimulator of interferon (IFN) genes (STING)-dependent pathogen mimicry response (PMR) in ovarian (OC) and other cancers. We now show that combining DNMTis and PARPis upregulates expression of a little-studied nucleic-acid sensor, NFX1-type zinc finger-containing 1 protein (ZNFX1). We demonstrate that ZNFX1 is a novel master regulator for PMR induction in mitochondria, serving as a gateway for STING-dependent PMR. In patient OC databases, high ZNFX1 expression levels correlate with advanced stage disease. ZNFX1 expression alone significantly correlates with an increase in overall survival in a phase 3 trial for therapy-resistant OC patients receiving bevacizumab in combination with chemotherapy. In correlative RNA-seq data, inflammasome signaling through ZNFX1 correlates with abnormal vasculogenesis. ZNFX1 controls PMR signaling through the mitochondria and may serve as a biomarker to facilitate offering personalized therapy in OC patients, highlighting the strong translational significance of our findings.

**Significance statement:** DNA methyltransferase and poly(ADP-ribose) polymerase inhibitors upregulate expression of a novel nucleic-acid sensor, ZNFX1 that serves as a mitochondrial gateway to STING-dependent interferon/inflammasome signaling with tumor suppressor properties in ovarian cancer.

## Introduction

Anti-cancer therapies that target epigenetic modulation, such as DNA methylation, induce inflammasome signaling, immune cell attraction and enhanced efficacy of immune checkpoint therapy (1–4). DNMTis 5-azacytidine (AZA) or decitabine (DAC) promote demethylation and transcription of endogenous retroviral (ERV) elements encoded within the eukaryotic genome, leading to accumulation of cytosolic dsRNA transcripts that induce type I IFN signaling (1). Moreover, combining DNMTi with a poly (ADP-ribose) polymerase inhibitor (PARPi) potentiates this effect by inducing type I IFN via DNA damage and stimulator of IFN genes (STING)-dependent cytosolic dsDNA sensor activation, a mechanism we term “pathogen mimicry response” or PMR (5,6).

A key gateway for inflammasome signaling is the mitochondria (mt)-mediated defense response (7,8). Mitochondrial antiviral signaling protein (MAVS), located in the mt outer membrane, is essential for RIGI-like helicases (RLH)-mediated antiviral signaling, activating both type I IFN transcription via the TBK1-IRF3/7 axis and inflammatory cytokine release via IKK-NFκB (8). The STING pathway, critical for mt involvement in the antiviral immune response (9), activates IFN and NFκB signaling through detection of dsDNA in the cytosol. Recent evidence shows that viral infection-induced mt dysfunction results in leakage of mtDNA into the cytosol and STING-dependent IFN and inflammasome pathway activation, serving as a key mediator of innate immune responses (10,11). However, the significance of this key process in cancer remains to be established.

A recent report by Vavassori *et al.* describes a rare autosomal recessive deletion of a little-studied protein, NFX1-type zinc finger-containing 1 protein (ZNFX1), which presents with severe immunodeficiency and multisystem inflammation following viral infection, often leading to death in early childhood (12). Upon viral infection, ZNFX1 shuttles from the cytosol to the mt outer membrane, where it binds viral dsRNA and interacts with MAVS to increase expression of interferon-stimulated genes (ISGs) (13). In this scenario, ZNFX1 acts as a very early, mt-dependent step for immune activation in defense against viruses (13), but how ZNFX1 mediates these processes is not well understood in general and especially not in cancer.

We now report a master-like role for ZNFX1 as a mediator of mt responses to the presence of dsRNA/dsDNA in the context of DNMTi and PARPi treatment. The combination treatment induces mtROS, mtDNA damage and subsequent release of damaged mtDNA into the cytosol, culminating in the induction of STING-dependent IFN and inflammasome signaling in ovarian cancer (OC) cells. CRISPR KO of ZNFX1 in OC cells ablates this signaling and reveals tumor suppressor gene (TSG)-like activity *in vitro* and *in vivo*. Importantly, we show that high ZNFX1 expression alone correlates with a significant increase in overall survival (OS) in patients with recurrent platinum-resistant OC receiving the angiogenesis inhibitor bevacizumab, providing the first key evidence for the translational potential of ZNFX1 in cancer.

## Materials and methods

### Bioinformatics analysis

Raw TCGA counts for ovarian serous cystadenocarcinoma (OC), triple-negative breast invasive carcinoma (TNBC), and colon adenocarcinoma (COAD) were obtained through Broad Institute GDAC Portal. These raw count data were processed using EdgeR and Limma powers differential expression analyses for RNA-sequencing and microarray studies. For the comparison between ZNFX1 high vs ZNFX1 low samples were split into respective groups based on median normalized count expression. Volcano plots were generated using pheatmap: Pretty Heatmaps. R package version1.0.12. https://CRAN.R-project.org/package=pheatmap. Pathways analysis was conducted using Clusterprofiler (14), an R package for comparing biological themes among gene clusters (15), tidyverse (16) and enrichplot (17). Manually curated dot plots were generated using ggplot2 (18).

Microarray expression data were extracted from the GEO using GEOquery (15), including datasets GSE9891 (19), GSE30161 (20), GSE26193 (21–24). The Z scores of gene expression levels were calculated according to normalization on all genes within samples then across all samples for the same data set. Raw counts of genes from the RNA-seq, GSE211669 (25,26) and GSE102118 (27), were downloaded from the GEO then converted to FPKMs using edgeR (28,29) followed by Z scores normalized in the same way as what for microarray expression data. Stage and grade information were retrieved from the GEO and published papers. Wilcoxon test was conducted to determine statistical significance of differences between stage and grade regarding ZNFX1 and CMPK2 expression. The correlations between ZNFX1 and CMPK2 expression were calculated as Pearson corrections with statistical significance. Scatter plots between ZNFX1 and CMPK2 expression were plotted using ggpubr R package (30).

We employed the pre-cancer atlas dataset (please refer to the biorxiv version of the paper: https://www.biorxiv.org/content/10.1101/2024.09.25.615007v1). In brief, we utilized micro-regional spatial whole transcriptome (GeoMx) (NanoString, Seattle, USA) dataset described in this study. The specimens were collected from both the incidental group (i.e. no cancer was diagnosed as a part of risk reduction surgeries or opportunistic salpingectomy) and cancer group. We only utilized the regions of the epithelial of the fallopian tube, fimbriae, p53 signature, Serous Tubal Intraepithelial Carcinoma (STIC), and cancer. STIC was collected from both incidental and cancer group. All sample processing and sequencing were performed by the Dana Farber Sequencing or HMS facility. The quality control (QC) and the Quartile-3 (Q3) normalization of the initial data set were performed as suggested by NanoString using GeoMx DSP software, NanoString (v 3.1.0.221). The details of the method and QC of the data will be found in the method and Supplementary Methods section of the pre-cancer atlas study.

### Pan-cancer bulk RNA-seq analysis

The bulk transcriptome profiles of ZNFX1 were downloaded from the TNMplot database which utilized datasets from NCBI GEO, GTex, TCGA, and TARGET databases (31). For each sample, the transcript read counts were normalized using MAS5 algorithm for NCBI GEO, GTex datasets and DESeq2 algorithm for TCGA and TARGET datasets to ensure uniform stability in the data.

### Cell lines

A2780 and TyKNu cells (a gift from Dr. Stephen Baylin) were cultured in RPMI-1640 (Corning) supplemented with 10% FBS (Sigma) and 1% penicillin-streptomycin (Sigma). OVCAR4 cells (a gift from Dr. Kenneth Nephew) were cultured in DMEM (Gibco) supplemented with 10% FBS, 1% penicillin-streptomycin, 1% Minimum Essential Medium vitamin solution (Corning) and 1% non-essential amino acids (Corning). The KPCA cell lines were developed and described in Iyer et al (32). Epithelial ovarian cancer cell lines (CP70, A2780, HeyC2, C272) were maintained in RPMI-1640 (Invitrogen, Carlsbad, CA) supplemented with 10% fetal bovine serum (FBS) (ATCC, Manassas, VA, USA) and 1% Penicillin-Streptomycin Solution (ATCC, Manassas, VA, USA). OVCAR8, OV2008, HEYA2 were maintained in DMEM (Invitrogen, Carlsbad, CA) supplemented with 10% fetal bovine serum (FBS) (ATCC, Manassas, VA, USA) and 1% Penicillin-Streptomycin Solution (ATCC, Manassas, VA, USA). To ensure cell line integrity, all cell lines were thawed at frequent intervals and not used beyond 40 passages. Additionally, cell morphology was monitored for each cell line, and proper media and growth conditions selected. All cell lines were cultured at 37LJC with 5% CO2. All cells were regularly tested for mycoplasma.

### In vitro treatments

5-Azacytidine (Sigma) was prepared at 500μM in PBS. Talazoparib (Biomarin) was prepared at 5mM in DMSO. Rotenone (Sigma) was prepared at 10μM in DMSO. In vitro treatments were performed as indicated in the text, with mock treatments performed using an equivalent final concentration of DMSO.

Transfection of the double-stranded DNA mimic polyinosinic: polycytidylic acid (poly I:C) or the double-stranded RNA mimic polydeoxyinosinic:deoxycytidylic acid (poly dI:dC) was performed using Lipofectamine 3000 (Invitrogen), followed by 72h incubation.

### Quantitative real-time PCR (qRT-PCR)

Total RNA was isolated after 3 or 6 days of treatment for qRT-PCR analysis to measure mRNA abundance of the indicated genes, normalized to GAPDH and β-actin mRNA **(Table1).** Data presented are the fold change after drug treatment over mock by ΔΔCT method.

**Table1.**
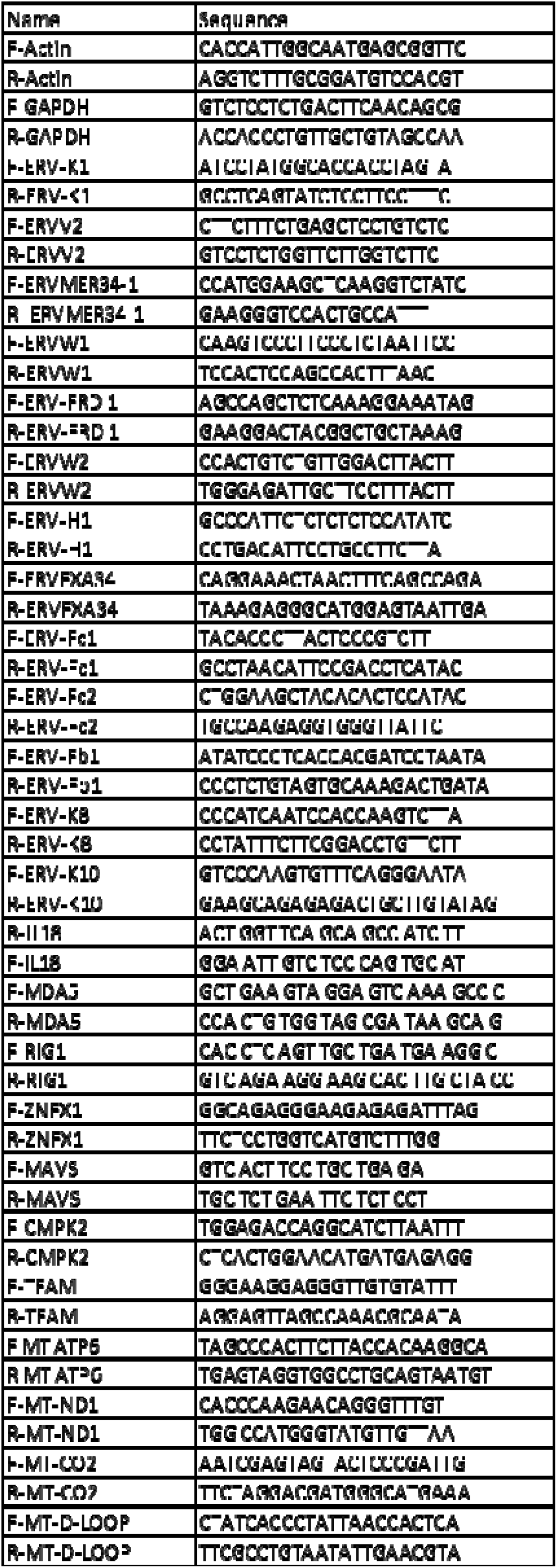

### Immunoblotting

Total cellular protein was extracted in RIPA buffer (Sigma) after 3 or 6 days of treatment. All gels were processed in the same way, In cases when the amount of samples were exciding the gel capacity, two gels were used side by side, similarly processed and normalized by the appropriate control. Mitochondrial protein fraction was isolated according to kit protocol (Mitochondrial Isolation Kit, Abcam). The following antibodies were employed to determine protein abundance: anti-ZNFX1 (1:1000, Abcam, #ab179452), anti-MAVS (1:1000, Abcam, #ab290729), anti-TFAM (1:1000, Cell Signaling, #8076), anti-VDAC (1:1000, Cell Signaling, #4866)., STAT3 (1:1000, Cell Signaling, #4904), NFKB 105/50 (1:1000, Cell Signaling, #12540), NFKB 65 (1:1000, Cell Signaling, #8242) pTBK1 (1:1000, Cell Signaling, #5483), TBK1 (1:5000, Abcam, #ab40676), STING (1:1000, Cell Signaling, #13647), pSTING (1:1000, Cell Signaling, #19781), Vinculine (1:1000, Cell Signaling, #13907)

### Immunofluorescence and proximity ligation assay

Treated cells were plated onto coverslips, fixed in 4% paraformaldehyde, and permeabilized in 0.1% triton x-100 in PBS.

For immunofluorescence, coverslips were blocked in 10% FBS in PBS, then incubated with primary antibody against pSTING (rabbit, 1:50, Cell Signaling) and anti-mouse Dylight 594 (ThermoFisher). Coverslips were mounted on slides using ProLong Gold Antifade Reagent with DAPI.

For proximity ligation assay, coverslips were blocked in 5% goat serum in PBS, then incubated with primary antibodies against ZNFX1 (rabbit, 1:50, Abcam), MAVS (mouse, 1:50, Invitrogen), or dsRNA (mouse, 1:50, EMD Millipore). Duolink in situ proximity ligation assay was performed per manufacturer protocols (Sigma), and coverslips were mounted on slides using ProLong Gold Antifade Reagent with DAPI.

Foci were examined using a Nikon Eclipse 80i fluorescence microscope (100×/1.4 oil, Melville, NY).

### Detection of reactive oxygen species

Flow cytometric detection of total cellular or mitochondrial ROS was performed following incubation of cell suspensions for 30min at 37°C with dihydroethidium (3μM, Invitrogen) or MitoSOX (3μM, Invitrogen), respectively.

### Cytosolic mtDNA detection

Cytosolic fractions were isolated (Mitochondria Isolation Kit for Cultured Cells, Thermo Scientific) and DNA was extracted (QIAamp DNA Blood Mini Kit) according to manufacturer protocols. qPCR examining cytosolic DNA content was performed using primers against mitochondrial (mt-ATP 6, mt-CO2, mt-ND1, D-loop) and nuclear (GAPDH, β-actin) genes (table 1). Relative cytosolic DNA quantity was normalized against total cellular GAPDH and β-actin mRNA isolated from the same input samples.

### Mitochondrial DNA damage detection by long PCR

Total cellular DNA was examined by quantitative PCR optimized for detection of ∼8-12kb fragments as previously described (33) using GoTaq Long PCR polymerase (Promega), Evagreen (1:20 Biotium), and 1x ROX reference dye (Invitrogen). Primers targeted an 8.9kb long mitochondrial fragment or 221bp short mitochondrial reference sequence (reference to primer table goes here). Cycling conditions for long fragment PCR: hot start 95°C 2min; denaturation 95°C 10s, extension 68°C 4min30s (x40 cycles). Cycling conditions for short fragment PCR: PCR: hot start 95°C 2min; denaturation 95°C 10s, extension 60°C 45s (x40 cycles); final extension 72°C 10min.

### Detection of cytokines release by ELISA

Cytokines release from the cells were measured in both WT cells and ZNFX1 KO cells in triplicates using ELISA kit according to manufacturer protocol. Cells were treated with different conc of AZA and Tal and ELISA was performed, absorbance was taken at 450 nm using a VersaMax ELISA Microplate Reader from Molecular devices. IFN-γ was detected by using Invitrogen™ Human IFN-γ ELISA kit and TNF-α was detected by using Invitrogen™ Human TNF-a ELISA kit.

### 8-hydroxy-2-deoxyguanosine determination using ELISA

mtDNA was isolated from both WT cells and ZNFX1 KO cells and 8-hydroxy-2-deoxyguanosine were detected using an ELISA kit as described in the manufacturer protocol. Cells were treated with different conc of AZA and TAL and ELISA was performed, absorbance was taken at 450 nm using a VersaMax ELISA Microplate Reader from Molecular devices. 8-hydroxy-2-deoxyguanosine was detected by using an abCam Human 8-hydroxy-2-deoxyguanosine ELISA kit.

### CRISPR CAS9 KO

CRISPR cell lines exhibiting genetic knockout of the ZNFX1 gene were generated in the Translational Laboratory Shared Services CRISPR Core (TLSS-CRISPR) using the CRISPR-Cas9 mechanism with synthetic single-guide RNAs (sgRNAs, Synthego) were generated targeting exons 6, 3, 8, and 11 (sequences below). CRISPR-Cas9 KOs were produced by nucleofection on the Lonza Amaxa™ 4D-Nucleofector platform and confirmed by subjecting cells to PCR and Sanger sequencing. Genomic editing was confirmed by INDEL analysis using the Synthego ICE analysis platform. Clonal KO populations were then generated by single-cell plating and screening of clonal sequences using ICE analysis.

sgRNA Sequences, Human:

Exon 6: ACCCTGGAGTGCACCATGCG

Exon 3: GGAGTGTAACTCTCATGTGA

Exon 8: GCCATGAGGCTAGACCATTG

Exon 11: GGTGGTCCCCAATCAAAATG

### Mitochondrial DNA transfection Assays

Total cellular DNA was purified from untreated cells by spin column extraction (Qiagen). Mitochondrial DNA was PCR amplified using a REPLI-g Mitochondrial DNA kit (Qiagen) and fragmented using DNAse I (New England Biolabs). Fragmented DNA was transfected into cells using Lipofectamine 3000 (Invitrogen). At 72h, cellular RNA was collected and assayed by qRT-PCR for expression of interferon-stimulated genes.

### Cell Doubling

Cells were seeded onto 24 well plates at a density of 50,000 cells/well on Day 0 and counted 3 wells at every 24 hours’ time point using hemocytometer. Cell number was quantified by plotting number of cells against period.

### Proliferation assay

Cells were seeded into 96-well plates at a density of 500 or 1000 cells/well and incubated for 2, 4, 6 days (TYK-nu) or 1, 2, 3 days (KPCA). Proliferation was assayed using MTS assay (Promega) according to the manufacturer’s instructions. Three replicate wells were used for each condition. Absorbance was measured at 490 nm using a spectrophotometer microplate reader.

### Transwell migration assay

The trans-well migration assay was previously described in (34). In short, Boyden chambers (8 μm pore size; Corning) were placed in the wells of a 24-well plate filled with 750 µl of serum-containing media which is used as a chemoattractant. 5.0 × 104 cells suspended in 500 µl of serum-free media then plated into a Boyden chamber and allowed to migrate for 16 hours. Following incubation, medium was aspirated from the Boyden chambers, the internal portion of the membrane was washed with phosphate-buffered saline (PBS) and cotton swabs, and the membrane was stained with Hema 3 staining kit. The membranes dried for 24 h before being plated on microscope slides. Each condition was performed in duplicate, five images were taken per membrane and cells were counted. Cells were imaged using the 5X objective and counted using ImageJ.

### Wound Healing

Briefly, cells were counted at a concentration of 5 × 10^5 cells/mL in cell culture media. The culture insert was aseptically placed at the bottom of a 12-well plate. Approximately 70 µl of cells were added to each side of the insert, and 1 ml of fresh media was introduced into the well surrounding the insert. The cells were allowed to reach a confluent monolayer over 12-24 hours in a 37-degree incubator. Prior to commencing the assay, verification of cell confluence inside the insert was conducted. After a 2-hour incubation, the insert was carefully removed to avoid disrupting the cell layer. Media was gently aspirated, and a 1 ml PBS rinse was performed. Fresh media containing 2% FBS was added gently to the side of the well to prevent cell detachment. Cell imaging using a microscope was carried out every 6 hours, capturing at least 2 images per well. For quantification of the gap area and calculation of the percentage change in area for each cell line or condition, an ImageJ plugin designed for high-throughput image analysis of in vitro scratch wound healing assays was utilized.

### Colony formation assay

TYK-nu and KPCA cells were plated at a concentration of 1,000 cells/well and 200 or 500 cells/well in a 6-well culture plate (Corning), with pre-warmed growth media. The cells were evenly dispersed by gently rotating the plate and then incubated for 7-10 days. Following the incubation, cells were fixed with 10% formalin and stained with 0.5% crystal violet in 25% methanol. The plates dried, and the colonies were imaged and counted with the Genesys software (Syngene). Each value reported is the mean of three biological replicates, each derived from the mean of three technical replicates (35).

### Spheroid assay

Cells were initially seeded at a 60–70% confluency level in 10cm plates. Subsequently, 3000 cells (TYK-nu) or 1000 cells (KPCA) were plated in triplicates in 24-well ultra-low adherent plates (Corning, cat #3473) with 1ml of stem cell medium, following a previously described protocol3. The cells were allowed to grow for 7-10 days.

Evaluation of spheroid number and area was conducted using a Zeiss Axiovert 40 inverted microscope equipped with Axio-Vision software (Carl Zeiss MicroImaging). Spheres or clusters smaller than 100 μm were excluded from the analysis (36).

### Cell cycle analysis

For cell cycle analyses, approximately 2.5×105 cells were plated in a 10 cm dish, allowed to attach overnight. Cells were harvested and fixed overnight in ice cold 70% ethanol and stored at 20oC until PI staining. Fixed cells were pelleted, washed in PBS, and incubated with RNase (0.1 mg/mL) at 37°C for 30 minutes. Cells were pelleted again, washed in PBS to remove the RNase, and then resuspended in PI stain solution (0.1 mg/mL) with a final cell concentration 1×106 cell/mL. The cells were then incubated on ice for 30 min and analyzed by LSRII flow cytometry analyzer with FACSDiva software.

### Immunohistochemistry

Tumors excised from mice were fixed overnight in 10% formalin, embedded in paraffin, and sectioned. The sections were stained with hematoxylin and eosin (HE) and photographed using a Leica light microscope at ×200 magnification. The sections underwent immunohistochemical staining using routine methods. Briefly, sections (5 μm) were deparaffinized, endogenous peroxidase was inactivated in 3% peroxide for 10 min, and antigen retrieval in 0.1 M sodium citrate was performed in a pressure cooker before the sections were blocked with 5% BSA and incubated overnight at 4 °C with polyclonal antibodies against ZNFX1, pSTING, pTBX1, WT1, PAX8, CD34. Primary antibodies were detected using SignalStain® Boost Detection Reagent (Rabbit: 8114, Mouse: 8125) and developed with SignalStain® DAB Substrate Kit followed by dehydration with increasing alcohol solutions and mounted. Slides were imaged with Motic EasyScan scanner and analyzed with QuPath software.

### Statistical Analysis

All data are presented as mean ± SEM with statistical significance derived from two-tailed unpaired Student’s t-test (or ANOVA).

The data generated in this study are publicly available in The Cancer Genome Atlas [TCGA], Gene Expression Omnibus [GEO] (including datasets GSE9891, GSE30161, GSE26193, GSE211669, GSE102118), Genotype-Tissue Expression (GTEx), and Therapeutically Applicable Research to Generate Effective Treatments [TARGET].

## Results

### ZNFX1 expression activates IFN and inflammasome signaling and is linked to mt dysfunction and a tumor suppressor signature

Our analysis of RNA-seq data from multiple Cancer databases (The Cancer Genome Atlas [TCGA], Gene Expression Omnibus [GEO], Genotype-Tissue Expression (GTEx), and Therapeutically Applicable Research to Generate Effective Treatments [TARGET] reveals that ZNFX1 expression is significantly altered in multiple cancers, compared with normal tissue counterparts, including, ovarian cancer (Figure 1A, S1). Genetic alterations in copy number are associated with changes in expression (Figure S1 B, C). ZNFX1 expression has a positive association with IFN/inflammasome genes mediating innate immune, type I IFN and dsDNA/RNA sensing (Figure 1B), but is inversely associated with expression of genes mediating mt function, including metabolism, in MSigDB (Figure 1C) (37) and MITOCARTA 3.0 (Figure 1D) (38) databases. Additionally, TCGA analyses in MSigDB (37) and MITOCARTA 3.0 (Figure 1D) (38) databases for triple-negative breast cancer (TNBC) (Figure S2A-C) and colon adenocarcinoma (COAD; Figure S3A-C) yields similar findings as above. Furthermore, CRISPR gene knockout (KO) of ZNFX1 in BRCA-proficient, high grade serous (HGS) OC cell line TYK-nu (Figure S4A) confirms ZNFX1 association with IFN/inflammasome genes and pathways, resulting in 604 downregulated and 443 upregulated differentially-expressed genes (Figure 1E). Additionally, gene-set enrichment analysis (GSEA) demonstrates suppression of IFN and inflammasome signaling by ZNFX1 KO (Figure 1E,F) and activation of pathways and expression of leading-edge genes involved in tumorigenesis, including, proliferation, migration and stemness (Figures 1E,F, S4B-E), which we explore in more detail below. *These data suggest that ZNFX1 and IFN/inflammasome signaling correlate with a mt dysfunction and tumor suppressor-like signature in OC cells*.

**Figure 1.**
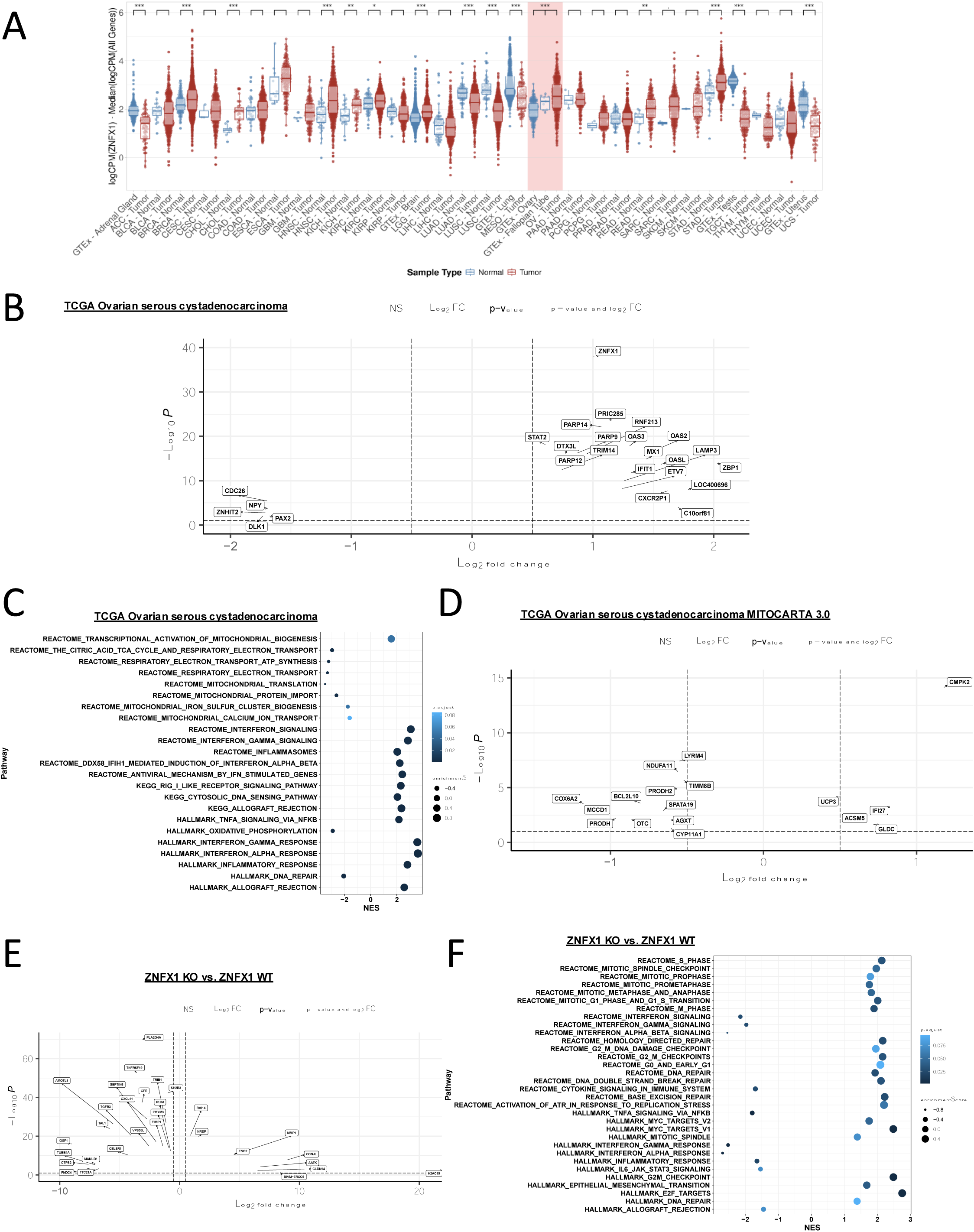
ZNFX1 expression correlates with interferon/inflammasome signaling, but is inverse to a mt dysfunction signature in OC cells. **A.** Pan-cancer analysis shows significantly higher relative expression of ZNFX1 in ovarian tumors vs fallopian tube normal samples. Raw RNA-seq expression counts for TCGA and GTEx samples were transformed to log2 counts-per-million values. The log ratio of ZNFX1 expression to the median expression of all genes in a sample is shown on the y-axis for each tissue type. A Wilcoxon Rank Sum test was performed on all tumor (shown in red) vs normal (shown in blue) samples within each tissue type. Unadjusted p-values (***p<0.001, **p<0.01, *p<0.05, . p<0.1) for each test are shown above each comparison. **B.** Volcano plot for RNAseq differential expression analysis (TCGA Ovarian serous cystadenocarcinoma), all annotated HGNC symbols, x-axis: log2 fold change in expression: ZNFX1 above median vs. ZNFX1 below median, y-axis: -log10 of FDR controlled adjusted p-value (padj), color mapping: gray: padj> 0.10 and log2 fold change < |0.5|, black: padj< 0.10 and log2 fold change < |0.5|, blue: padj> 0.10 and log2 fold change > |0.5|, and orange: padj< 0.10 and log2 fold change > |0.5|. **C.** Pathway dot plot depicting result from gene set enrichment analysis (TCGA Ovarian serous cystadenocarcinoma) on pre-ranked gene list derived from ZNFX1 above median vs. ZNFX1 below median differential expression analysis. Pathways depicted are derived from manual curation of Interferon, Mitochondria, and DNA repair pathways compiled from MSigDB: HALLMARK, KEGG, and REACTOME. x-axis: normalized enrichment score, dot size: enrichment score, color gradation: FDR controlled adjusted p-value. **D.** Volcano plot for RNAseq differential expression analysis (TCGA Ovarian serous cystadenocarcinoma), MITOCARTA 3.0 symbols, x-axis: log2 fold change in expression: ZNFX1 above median vs. ZNFX1 below median, y-axis: -log10 of FDR controlled adjusted p-value (padj), color mapping: gray: padj> 0.10 and log2 fold change < |0.5|, black: padj< 0.10 and log2 fold change < |0.5|, blue: padj> 0.10 and log2 fold change > |0.5|, and orange: padj< 0.10 and log2 fold change > |0.5|. **E.** Volcano plot for RNAseq differential expression analysis (ZNFX1 KO vs. ZNFX1 WT), all annotated HGNC symbols, x-axis: log2 fold change in expression, y-axis: -log10 of FDR controlled adjusted p-value (padj), color mapping: gray: padj> 0.10 and log2 fold change < |0.5|, black: padj< 0.10 and log2 fold change < |0.5|, blue: padj> 0.10 and log2 fold change > |0.5|, and orange: padj< 0.10 and log2 fold change > |0.5|. **F.** Pathway dot plot depicting result gene set enrichment analysis on pre-ranked gene list derived from ZNFX1 KO vs ZNFX1 WT RNAseq comparison. Pathways depicted are derived from manual curation of Interferon, Mitochondria, and DNA repair-associated pathways compiled from MSigDB: HALLMARK, KEGG, and REACTOME. x-axis: normalized enrichment score, dot size: enrichment score, color gradation: FDR controlled adjusted p-value.

### ZNFX1 is required for MAVS localization, mt dysfunction and dsDNA leakage into the cytosol

To expand on the hypothesized role of ZNFX1 as an early PMR defense mechanism for immune activation via mt-mediated mechanisms, we investigated whether ZNFX1 not only binds cytosolic nucleic acids but is crucial for localization of MAVS to the mt membrane. For these studies, we first show robust ZNFX1 expression in HGSOC in the Cancer Cell Line Encyclopedia (CCLE) database (Figure S5A), we validate these studies by using RT-qPCR and western blot analyses (Figure S5B,C) and perform functional assays in TYK-nu, OVCAR4 and A2780 cells. We further show that transfection of synthetic dsRNA and dsDNA mimics (poly I:C [PI:C] and poly[dI:dC], respectively) increases expression of ZNFX1 (Figure S6A,B). Moreover, treatment with the DNMTi azacytidine (AZA) alone or in combination with PARPi talazoparib (TAL) mimics this PMR defense mechanism by increasing ERV transcription in OC cells (Figure S6C), in agreement with our previous results in other cancer types (1, 3–5), and associates with an increase in ZNFX1 expression (Figures 2A, S6D). AZA and TAL alone or the AZA-TAL combination increase levels of MAVS (Figure 2A, S6D) and colocalization of ZNFX1 to dsRNA (Figure 2B, S6E), dsDNA (Figure 2C, S5F) and MAVS (Figures 2D, S6G,H), as analyzed by proximity ligation (PLA) and immunofluorescence (IF) assays. Importantly, ZNFX1 KO slightly increases MAVS expression with TAL and combination drug treatment, but inhibits MAVS colocalization to mt membrane protein TOM20 (Figure 2A, E). Thus, we demonstrate that *ZNFX1 binds both dsRNA and dsDNA and plays a crucial role in MAVS localization to the mt outer membrane*.

**Figure 2.**
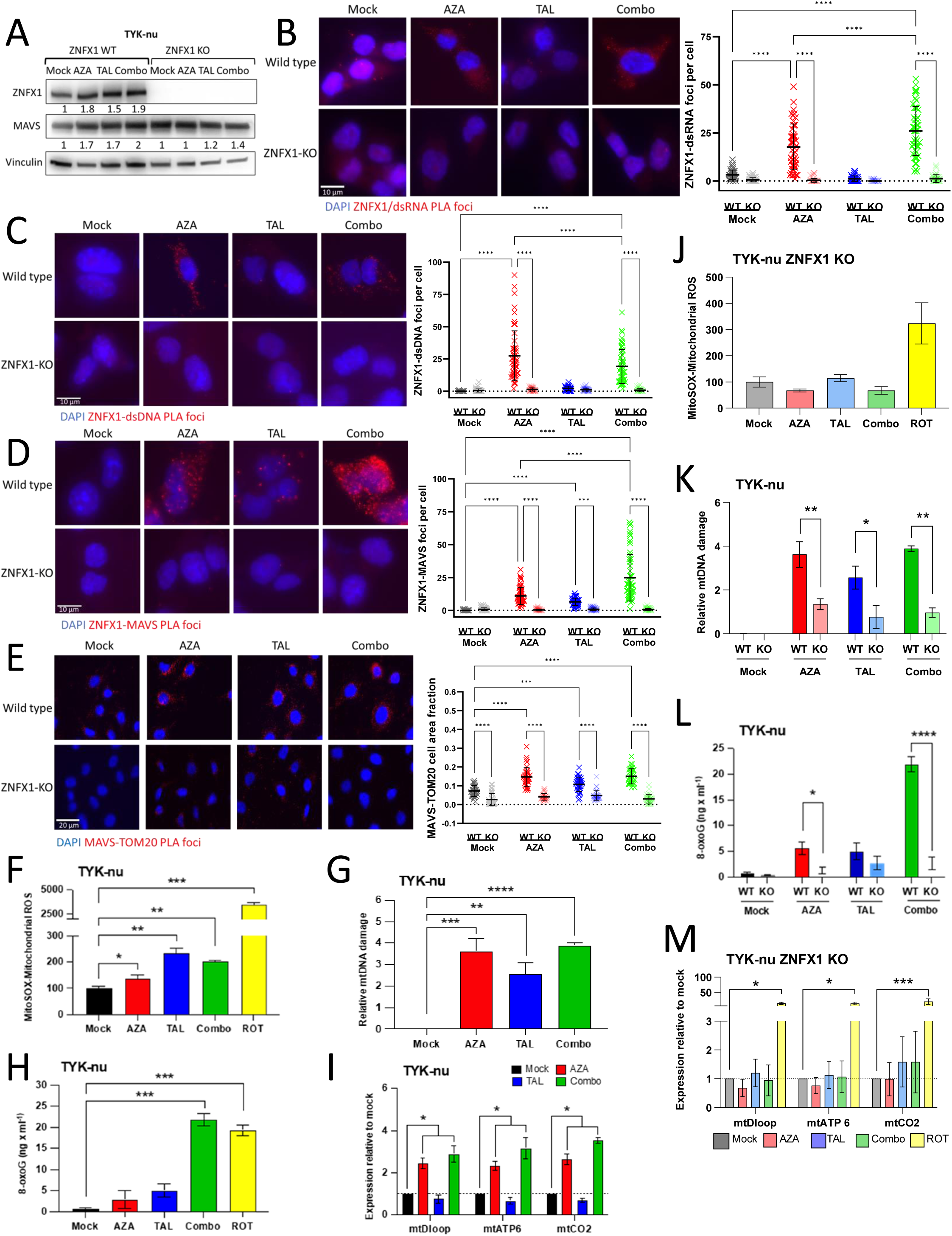
DNMTi and PARPi increase ZNFX1 expression, localization with MAVs, increasing mtROS, DNA damage, and dsDNA leakage into the cytosol. The following assays were performed in TYK-nu OC cells following 6 days of AZA 100nM, TAL 2.5nM, or combination treatment: **A.** Immunoblotting for ZNFX1 and MAVS. **B-E.** Representative immunofluorescence images of ZNFX1 interaction with dsRNA (C), dsDNA (D) MAVS (E), and MAVS and mt membrane protein TOM20 interaction (F), by proximity ligation assay (left): graphical representation of foci in three independent experiments is plotted (right). **F.** Flow cytometry detection of Mitosox measuring mtROS. **G.** Relative mtDNA damage measured by adapted real-time long-range PCR method. **H.** Relative 8-oxoG in mtDNA measured by ELISA. **I.** Relative expression of mt-encoded genes (mtDloop, mtATP6/8, mtCO2) in cytosolic DNA fractions quantitated by qPCR. **J-M.** The following assays were performed in ZNFX1 KO and/or WT TYK-nu OC cells following 6 days of AZA 100nM, TAL 2.5nM, or combination treatment: (J) Flow cytometry detection of mtROS in ZNFX1 KO TYK-nu cells following 6 days treatment with AZA, TAL, or combination. (K) Relative mtDNA damage measured by adapted real-time long-range PCR method in ZNFX1 WT and KO TYK-nu. (L) Relative 8-oxoG in mtDNA isolated from ZNFX1 KO and WT TYK-nu cells. (M) Relative expression of mt-encoded genes (mtDloop, mtATP6/8, mtCO2) in cytosolic fraction isolated from ZNFX1 KO TYK-nu. Rotenone used as a positive control in F, H, J, I and M. All data are presented as mean ± SEM with p values derived from two-tailed unpaired Student’s t-test or ANOVA as appropriate. * p<0.05, ** p<0.01, *** p<0.001, **** p<0.0001. All experiments were performed at least 3 times.

Given that ZNFX1 expression is required for MAVS localization, we hypothesized that ZNFX1 also plays a role in mt dysfunction, as measured by mt ROS (11), mt DNA damage and mt DNA leakage into the cytosol. Transfection of dsDNA/RNA mimics or treatment with AZA, TAL, or AZA-TAL combination markedly increase both mtROS levels (Figures 2F, S7A, B, C; flow cytometry analysis of specific mtROS dye, MitoSOX) and cellular ROS (35) (Figure S7C; dihydroethidium flow cytometry analysis) in OC cells (39). Moreover, using long-range PCR for targeting an 8.9kb long mt fragment for mtDNA damage as well as ELISA for levels of mt 8-oxoguanine (8-oxoG), we show that the above drug treatments increase mtDNA damage (Figures 2G,H, S7D-F). Finally, PCR analysis for mtDNA in cytosolic extracts demonstrates leakage of mtDNA into the cytosol due to AZA, TAL, or AZA-TAL combination treatments (Figures. 2I, S7G, H). Importantly, ZNFX1 KO (Figure 2A) abrogates each of these above steps (Figure 2J-M). *Taken together, these result support a master role for ZNFX1 in mt dysfunction*.

### ZNFX1 mediates DNMTi/PARPi-induced STING-dependent IFN/inflammasome signaling

The above dynamics suggest a candidate role for ZNFX1 in AZA and AZA-TAL-induced STING-PMR signaling (5). Accordingly, we show that these treatments (either for 3 or 6 days) induce ZNFX1-dependent increases in IFN/inflammasome gene transcripts and proteins, including TNFα, IFI27, MX2, CCL5 and CXCL10, and these increases are abrogated by knocking out ZNFX1 (Figure 3A, B, S8A-D). Additionally, forced expression of ZNFX1 (Figure S9A-C, 10A) rescues IFN/inflammasome signaling (Figure 3C,D, S10B). Likewise, AZA, TAL, or combination treatment increases levels of STING Ser366 phosphorylation (pSTING, i.e., active STING), as well as downstream pSTING targets pTBK1 and pIRF3 in TYK-nu, OVCAR4 and A2780 cells, and ZNFX1 KO abrogates these increases (Figure 3E, F, Figure S11A-C). KO of STING has the same effect as ZNFX1 KO (Figure 3G, H, S11D,E), confirming effects on STING pathway signaling. The role of mt mediation in the above dynamics is apparent following mtDNA transfection into ZNFX1 WT cells, leading to robust induction of TNFα, NFκB, IFI27, ISG15, and STING compared to untransfected controls (Figures 3I, S11F). Furthermore, ZNFX1 KO abrogates these changes (Figure 3I, S11F). Taken together, these data support *a master role for ZNFX1 in mediating mtDNA induction of STING-dependent IFN/inflammasome signaling*.

**Figure 3.**
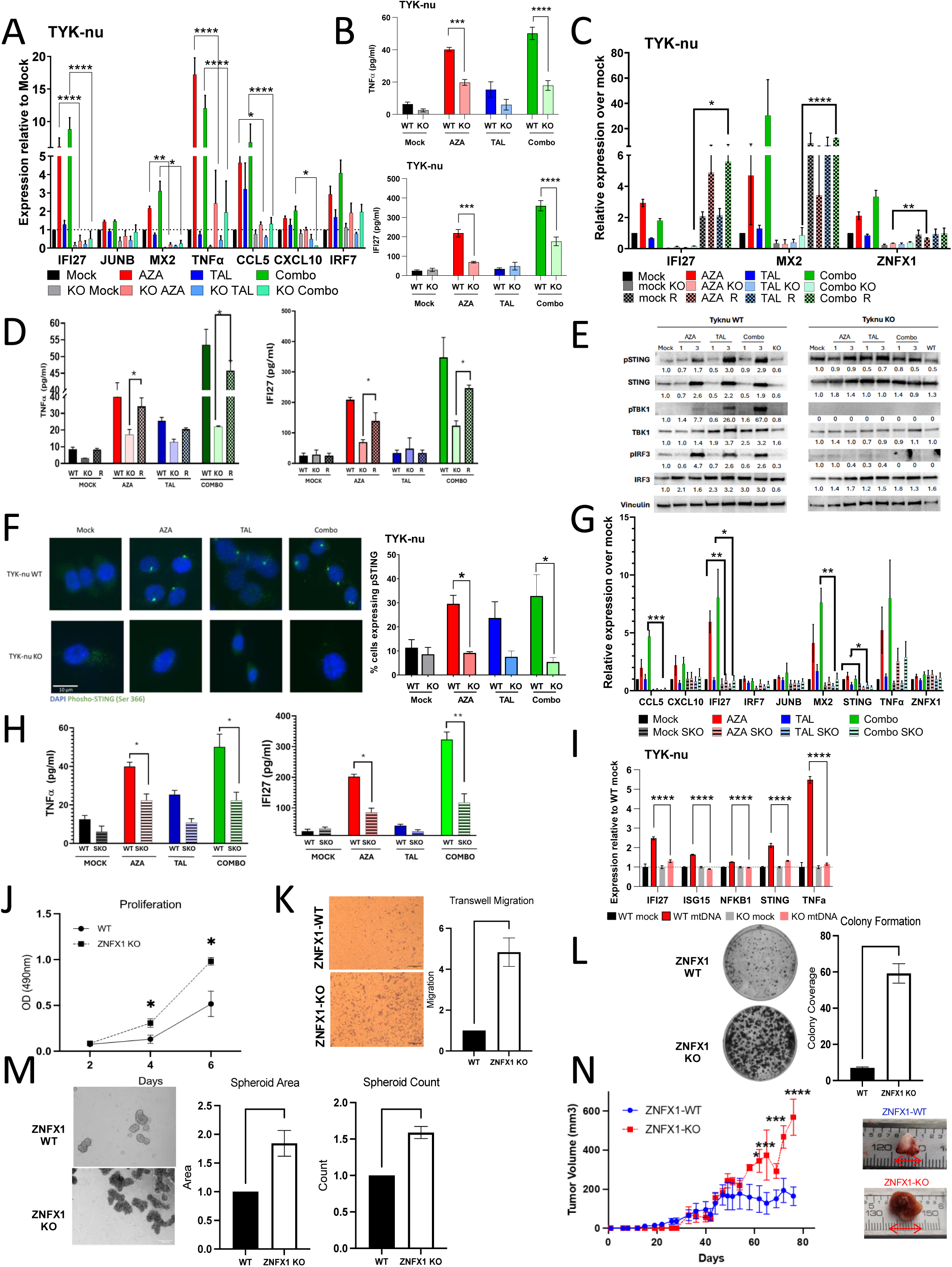
ZNFX1 increases DNMTi/PARPi-induced STING-dependent IFN and inflammasome signaling and ZNFX1 KO increases tumorigenic features in vitro and in vivo. **A.** Relative transcript levels of IFN (IFI27, MX2) or inflammasome (JUNB, TNFα) by qPCR in TYK-nu ZNFX1 WT or KO following 6 days treatment with 100nM AZA, 2.5nM TAL, or combination. **B.** Levels of cytokines, , TNFα (Top panel), IFI27 (Bottom panel), as measured by ELISA in TYK-nu ZNFX1 WT or KO following 6 days treatment with AZA 100nM, TAL 2.5nM, or combination in TYK-nu. **C.** Relative transcript levels of IFN (IFI27, MX2) or inflammasome (JUNB, TNFα) by qPCR in TYK-nu ZNFX1 KO cells transfected with ZNFX1 plasmid construct following 6 days treatment with 100nM AZA, 2.5nM TAL, or combination. **D**. Levels of cytokines, TNFα (Left panel), IFI27 (Right panel), as measured by ELISA in TYK-nu ZNFX1 KO cells transfected with ZNFX1 plasmid construct following 6 days treatment with AZA 100nM, TAL 2.5nM, or combination in TYK-nu. **E**. Relative expression levels of pSTING/STING, pTBKI/TBKI and pIRF3/IRF3 in protein extracts after 1 and 3 days of treatment with AZA, TAL and combination in TYK-nu. **F.** Representative immunofluorescence images of Ser366 phosphorylation of STING in TYK-nu ZNFX1 WT or KO following 24h treatment with AZA, TAL, or combination. **G**. Relative transcript levels of STING, IFN (IFI27, MX2, CCL5) or inflammasome (JUNB, TNFα) by qPCR in TYK-nu STING KO cells following 6 days treatment with 100nM AZA, 2.5nM TAL, or combination. **H.** Levels of cytokines, TNFα (Left panel), IFI27 (Right panel), as measured by ELISA in TYK-nu ZNFX1 STING KO following 6 days treatment with AZA 100nM, TAL 2.5nM, or combination in TYK-nu. **I.** Relative expression of IFN/inflammasome (IFI27, ISG15, NFKB1, STING, TNFα) transcripts by qPCR in TYK-nu ZNFX1 WT or KO 72hrs after transfection of purified mtDNA. **J-N.** Effect of ZNFX1 KO on TYK-nu (**J**) proliferation (seeding density 500 cells) **(K)** migration **(L)** colony formation (1000 cells/ well) **(M)** spheroid formation (3000 cells/well; 7-10 day growth period). and **(N)** tumor growth (3×106 TYK-nu WT or ZNFX1 KO cells injected S.C.; n=5 NSG mice per group) All data are presented as mean +/- SEM with p-values derived from two-tailed unpaired Student’s t test or ANOVA as appropriate. * p<0.05, ** p<0.01, *** p<0.001, **** p<0.0001. All experiments were performed at least 3 times.

### ZNFX1 KO increases tumorigenic features in vitro and in vivo

Our RNA-seq data from ZNFX1 KO TYK-nu cells (Figures 1D,E) shows activation of pathways involved in tumorigenesis, including hedgehog signaling pathway genes frizzled class receptor 4 (FZD4) and smoothened (SMO) (Figure 1E). These data prompted exploring functional analysis of tumorigenesis in ZNFX1 KO cells in human TYK-nu (Figure 12A-E) and/or mouse KPCA BRCA-proficient HGSOC cells (Figure S13A-J) (32). ZNFX1 KO increases proliferation and decreases cell doubling time (Figures 3J, S12A,B, S13B-C). Moreover, increases are seen in rate of wound healing (Figures S12C, S13D) and migration (Figures 3K, S13E), as well as colony (Figures 3L, S13F) and spheroid (Figures 3M, S13G) formation, with no effect on cell cycle dynamics (Figures S12D, S13H). Importantly, ZNFX1 KO increases tumor growth in xenograft mouse assays (Figures 3N, S12E). Forced expression of ZNFX1 (Figures S9A-C, 10A) rescues colony formation (Figure S13J). Knocking out STING induces similar changes in TYK-nu cells (Figures S14A-G). Immunohistochemical analysis of the tumors confirms loss of ZNFX1 in the KO tumors (Figure S15A, B) and a trend toward a decrease in p-STING (Figure S15A). In addition, ZNFX1-KO tumors show increased expression of epithelial tumor markers WT1 and CD31, expressed on early and mature vascular endothelial cells (Figure S15B). *Collectively, in the context of complete deletion, these functional assays strongly support a tumor suppressor-like (TSG) role for ZNFX1 in cancer*.

### Translational significance of ZNFX1 expression

To begin to investigate the translational significance of ZNFX1 in HGSOC, we examined clinical gene expression datasets (CSIOVDB (40) and TCGA). This analysis reveals potentially significant translational findings. *First*, ZNFX1 gene expression increases in precursor lesions of HGSOC found in the fallopian tube epithelium (STIC, Serous Tubal Intraepithelial Carcinoma) (Figure 4A)(41) and with increase tumor stage and grade (Figure 4B). Importantly, in the CSIOVDB dataset of 86 HGSOC patients, high ZNFX1 expression significantly correlates with increased overall survival (OS; p=0.032) (Figure 4C). *Second*, in support of DNMTis increasing ZNFX1 expression in HGSOC cell lines (Figure 2A), we queried RNA-seq data for pre- and post-treatment tumor biopsies in a phase II clinical trial testing a DNMTi with immune check point therapy in HGSOC patients (42). In 9 available paired samples (pre-treatment baseline Cycle 1 Day 1 versus post-treatment Cycle 2 Day 8), an increase in ZNFX1 expression is seen in 6 of the 9 patients (p=0.027) (Figure 4D). While n-values for trial outcomes in this small trial allow only a case match to ZNFX1 levels, several interesting relationships emerge: 1) Overall, pre-treatment ZNFX1 levels are higher in the 4 patients with responses (RECIST) than the other patients (Figure 4E); 2) post-DAC treatment, these values remain higher and increase further in 2 of the 4 patients (Figure 4E); and 3) deconvolution studies of the above bulk RNA-seq data demonstrates that when comparing post-vs pre-DAC treatment, the highest vs lowest ZNFX1 quartile levels correlate with statistically significant changes in key immune cell types (CD8, CD4, plasma B cells; Figure S16A-C), in keeping with improved immune response in some patients in the trial (42).

**Figure 4.**
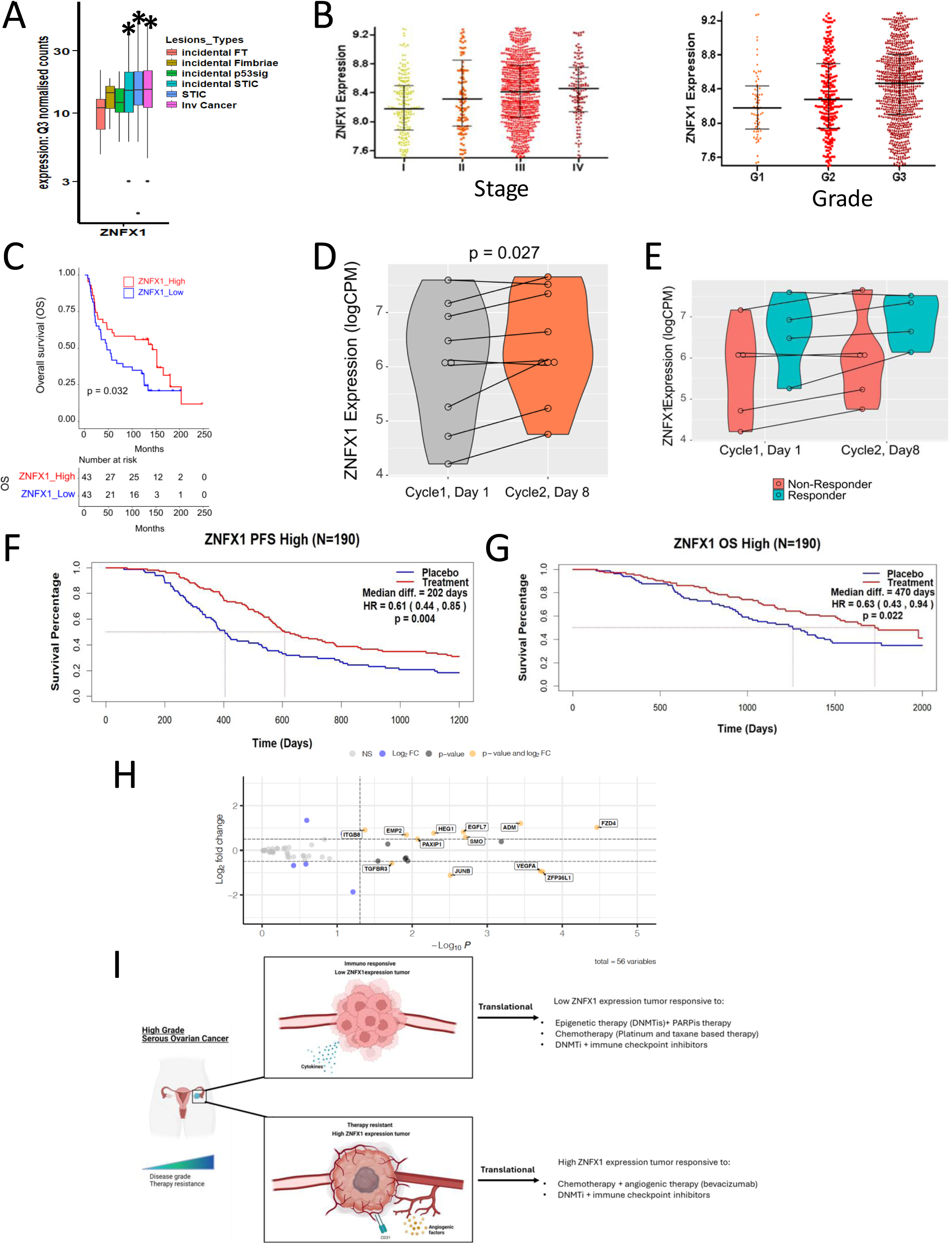
Translational relevance of ZNFX1 expression in ovarian cancer. **A.** This box plot depicts the Q3-normalised expression of ZNFX1 in the epithelia with HGSOC progression. The region of interest (ROI) for each lesion type in X axis was taken from micro regional spatial whole transcriptome (GeoMx). Number of ROIs per lesion type as follows: incidental FT (n=29), Incidental Fimbriae (n=26), incidental p53 signature (n=39), incidental STIC (n=27), STIC (n=96) and inv cancer (n=105). STIC = STIC associated with cancer and inv cancer = invasive HGSOC. Y axis is presented in log10 scale. The solid line indicates the median within the interquartile range, with whiskers extending to a maximum of 1.5 times the interquartile range beyond the box. Black asterisks indicate significant differences in stages compared to the FT.I; *p<0.05, Generalized Linear Mixed Models (GLMMs) taking patient ID as random effect**. B.** Expression of ZNFX1 in ovarian cancer patients in precursor lesions at different tumor stages (left) and grades (right) in the Ovarian Cancer Database of Cancer Science Institute Singapore (CSIOVDB). **C.** Overall survival plotted in CSIOVDB. **D.** Analysis of GSE188249 RNA-seq data (42) for ZNFX1 expression in 9 paired samples pre cycle 1 Day 1 (C1D1) and post Cycle 2 Day 8 (C2D8) epigenetic therapy. **E.** Analysis of RNA-seq data for ZNFX1 expression in samples from responders and non-responders pre (C1D1) and post (C2D8) epigenetic therapy. **F,G.** Kaplan-Meier curves progression-free survival and overall survival in ICON7 trial (standard treatment + bevacizumab v. standard treatment). High v. low ZNFX1 expression separated by median. **H.** Volcano plot for RNAseq differential expression analysis of curated vasculogenesis genes from TCGA: ZNFX1 above median vs. ZNFX1 below median, y-axis: -log10 of FDR controlled adjusted p-value (padj), color mapping: gray: padj> 0.10 and log2 fold change < |0.5|, black: padj< 0.10 and log2 fold change < |0.5|, blue: padj> 0.10 and log2 fold change > |0.5|, and orange: padj< 0.10 and log2 fold change > |0.5|. **I.** Graphical abstract showing effects of different therapies on basal levels of ZNFX1 as well as tumor responses. *Top panel:* High grade serous ovarian cancer cells with low basal levels of ZNFX1, DNMTI and PARPi (Figures 2-3), DNMTi and immune checkpoint inhibitors (Figures 4E, F) or chemotherapy as in the ICON7 trial (SFigure S14) can lead to tumor responses. *Bottom panel:* High grade serous ovarian cancer cells with high ZNFX1 expression and disease grade therapy resistance (ICON7 trial data, Figure 4G), cells may also exhibit immune evasive and angiogenic features that may contribute to responses to chemotherapy plus bevacizumab.

Finally, and of highest translational significance, we discover that high ZNFX1 expression correlates with outcomes in the ICON7 (43,44) phase III clinical trial, which tested the anti-VEGF drug bevacizumab added to platinum-based chemotherapy in patients with chemotherapy resistance. This 2011 trial saw an initial increase in progression-free survival (PFS, 3.8 month), but with no improvement in OS. This treatment paradigm (45) is not currently standard treatment in the US despite FDA approval (45). Our analysis of DASL (cDNA-mediated annealing, selection, extension and ligation) gene expression data from a subset of the ICON7 trial (43,44) now shows that low ZNFX1 expression correlates with significant (p<0.002) improvement of PFS in response to chemotherapy (SF 12), but not with OS (p value = 0.12 (SF12); Figure S17A, B). However, when combined with bevacizumab, high ZNFX1 expression not only significantly correlates with improvement in PFS of 6.6 months, but also with a marked significant improvement in OS of 15.6 months (Figure 4F,G). To shed light on the above ICON7 findings, our query of the TCGA database for gene expression changes associated with high ZNFX1 gene expression shows that abnormal vasculogenesis tracks with high ZNFX1 expression (Figure 4H). Of particular note, expression of CEACAM1 (carcinoembryonic antigen-related cell adhesion molecule) is a key leading edge gene change. This immune-inflammasome IRF1-driven gene, when over-expressed in the tumor microenvironment (TME), drives abnormal angiogenesis and immune T cell tolerance (Figure 4H) (46). Of note, in terms of potential control by ZNFX1 of the above signature, it is reversed in ZNFX1 KO cells with CEACAM1 now the most decreased leading-edge gene (Figure S4E).

Overall, our above data suggests that: 1) Increased ZNFX1 expression correlates with increased survival of HGSOC patients; 2) DNMTis increase ZNFX1 expression in a clinical setting, with distinct changes in key immune cell subsets; 3) High ZNFX1 expression in tumors from HGSOC patients with chemotherapy resistance and accompanying abnormal TME vasculature helps explain why treatment with the anti-angiogenesis drug bevacizumab correlate for the first time with significantly increased OS in a subset of ICON7 trial patients (see summary of these points in graphical abstract Figure 4I).

## Discussion

Our fundamental finding is that ZNFX1 plays a master regulator role in inducing mt-mediated STING-dependent IFN/inflammasome signaling in OC cells. In this paradigm, ZNFX1 is central to MAVS localization and induction of mt dysfunction, previously shown to be important for innate immune responses (7). Our study provides key new insights for how DNMTis and PARPis induce PMR (1–3,5) and their potential clinical impact as anti-cancer therapies (47).

Our data indicate that ZNFX1 is activated not only by dsRNA as previously reported (13), but also by dsDNA as well as viral mimics DNMTis and PARPis, culminating in mt dysfunction and leakage of mtDNA into the cytosol, resulting in activation of STING-dependent interferon and inflammasome signaling. In validation, we also show that transfecting mtDNA in OC cells activates STING signaling, while the same effect was not observed in ZNFX1 KO cells. Therefore, ZNFX1 could also potentially be mechanistically required for STING activation, suggesting future study of its interaction with STING is warranted. Another mechanism of activating ZNFX1 through mtDNA could potentially involve direct activation of cGAS (48) in the cytosol as the graphic summary suggests (Fig. 4I, S18). Furthermore, type I IFNs (IFN-I), dependent on Jak-STAT signaling (13) could also contribute to transcriptional regulation and activation of ZNFX1 in the context of the present study. Regardless of the mechanism involved, once in the cytosol, mtDNA will be detected by ZNFX1 in the similar manner as nuclear DNA and RNA and lead to STING activation.

Our studies also demonstrate that ZNFX1 also acts to suppress cell growth and neoplastic behavior, with tumor suppressor-like properties. Thus, when ZNFX1 is knocked out in vitro and in vivo in human and mouse OC cells, multiple tumorigenic phenotypes emerge. In contrast, when ZNFX1 is chronically expressed at high levels in therapy-resistant cancer cells, anti-tumor inflammasome signaling can lead to activation of vasculogenesis-induced immune evasion to enable cancer cell survival. It is now well established that tumors resurrect an embryonic vascular program to escape immunity (49). Blocking such effects in therapy scenarios, as is evident in our trial data for adding bevacizumab to chemotherapy (43,44), warrant studying the role of ZNFX1 in future basic, clinical and translational cancer biology investigations.

Our above data has high translational significance in the context of ZNFX1 expression in patients with HGSOC as follows: **1**) High ZNFX1 expression tracks with IFN/inflammasome immune signatures in primary HGSOC (TCGA and clinical trial datasets). ZNFX1 levels increase with increasing stage and grade of disease, and correlate with overall therapy response (discussed below); **2**) The DNMTi class of epigenetic drugs increase ZNFX1 in pre-clinical studies, which correlates with known induction of potent immune functions in the TME (47). We now have early *in vitro* evidence of efficacy in HGSOC, which needs rapid translation into a patient-based context to determine potential synergistic response with immune checkpoint therapies (42); **3**) Perhaps most important, ZNFX1 expression is a potentially robust independent biomarker for predicting therapy responses, per our analysis of in a large phase III trial in HGSOC. The majority of HGSOC patients develop recurrent, chemoresistant disease, limiting 5 year survival, and reversing this resistance is a great unmet need (50). Although the addition of bevacizumab to chemotherapy has shown promise by extending PFS in the ICON7 phase III trial and also the GOG218 trial, lack of durability and benefit for OS has prevented this therapy combination from gaining traction in routine HGSOC therapy, despite FDA approval (45). Our finding that high ZNFX1 expression tracks with an impressive increase in median OS of 15.6 months (Figure 4G), if further validated, suggests that ZNFX1 levels should be considered to personalize the use of bevacizumab in OC treatment.

In conclusion, our present findings reveal novel mechanistic and translationally significant roles for the little-studied gene, ZNFX1. Our studies demonstrate multiple complex properties of ZNFX1 that suggest a master role for controlling mt dynamics, resulting in inflammasome signaling responses to DNMTis and PARPis via STING-dependent-IFN/inflammasome induction (1–3,5,6). Translationally, levels of ZNFX1 may balance between tumor suppressor functions and immune functions linked to vascular integrity (49). The latter scenario reveals an important biomarker role for ZNFX1 levels predicting overall survival for HGSOC patients receiving bevacizumab therapy.

## Acknowledgements

Our studies were supported by the Adelson Medical Research Foundation (L.S., R.A., M.J.T., K.T., F.V.R., and S.B.B.); National Cancer Institute–Cancer Center Support Grant P30 CA134274 University of Maryland Marlene and Stewart Greenebaum Comprehensive Cancer Center (F.V.R.); NIEHS grant (2R01ES011858) (R.A., F.V.R., S.B.B.); NCI grant (1RO1CA259635-01A1) and 1RO1AG078814 (D.C.W), the Human Genetics Graduate Program, University of Maryland (L.S.); the Maryland Department of Health’s Cigarette Restitution Fund Program (F.V.R.); Specialized Program of Research Excellence (SPORE) program, through National Cancer Institute (NCI) grant P50CA254897 (L.S., R.A., K.T., F.V.R, S.B.B, M.J. T., J-P.I., K.P.N); and the Van

Andel Research Institute – Stand Up to Cancer Epigenetics Dream Team (S.B.B., F.V.R., K.P.N.). Stand Up To Cancer is a program of the Entertainment Industry Foundation, administered by the AACR. M.J.T. is a recipient of The Evelyn Grollman Glick Scholar Award. T.K, S.S, and R.D were funded by Gray Foundation and SPORE grant (P50CA228991).The content is solely the responsibility of the authors and does not necessarily represent the official views of the NIH. We thank Dr. Rena Lapidus and Brandon Cooper (UMGCCC, Translational Laboratory Shared Services [TLSS]) for CRISPR KO studies. We also thank Dr. Aksinijah Kogan for help with R script and bioinformatics early in the project. We thank Dr. Daniela Matei (Northwestern University). With gratitude to the Jerry and Peggy Throgmartin Family for their philanthropic investments in our work.

## Authorship Contributions

L.S., R.A., M.J.T., K.T., S.B.B., K.P.N., and F.V.R. designed research; L.S, R.A., K.T., C.M.C., S.R., E.A.F, G.H., S.L., J.W. and R.C. performed research and analyzed data; S.F., F. H., R.M., Z.G.,G.E.K., G.E.S. contributed to generation of original clinical trial data sets, and R.A., L.S., K.T., J.W.G., D.C.W, B.J.N.W., T.K, S.S, R.D, S.K., J.P., M.M.A.I, V.K.T.,G.C., J-P.I., K.D.M., M.J.T., S.B.B., K.P.N, and and F.V.R. wrote the paper.

