## Supplemental Figures for "ZNFX1 is a Novel Master Regulator in Epigenetically-induced Pathogen Mimicry and Inflammasome Signaling in Cancer"

### Supplemental Figure S1

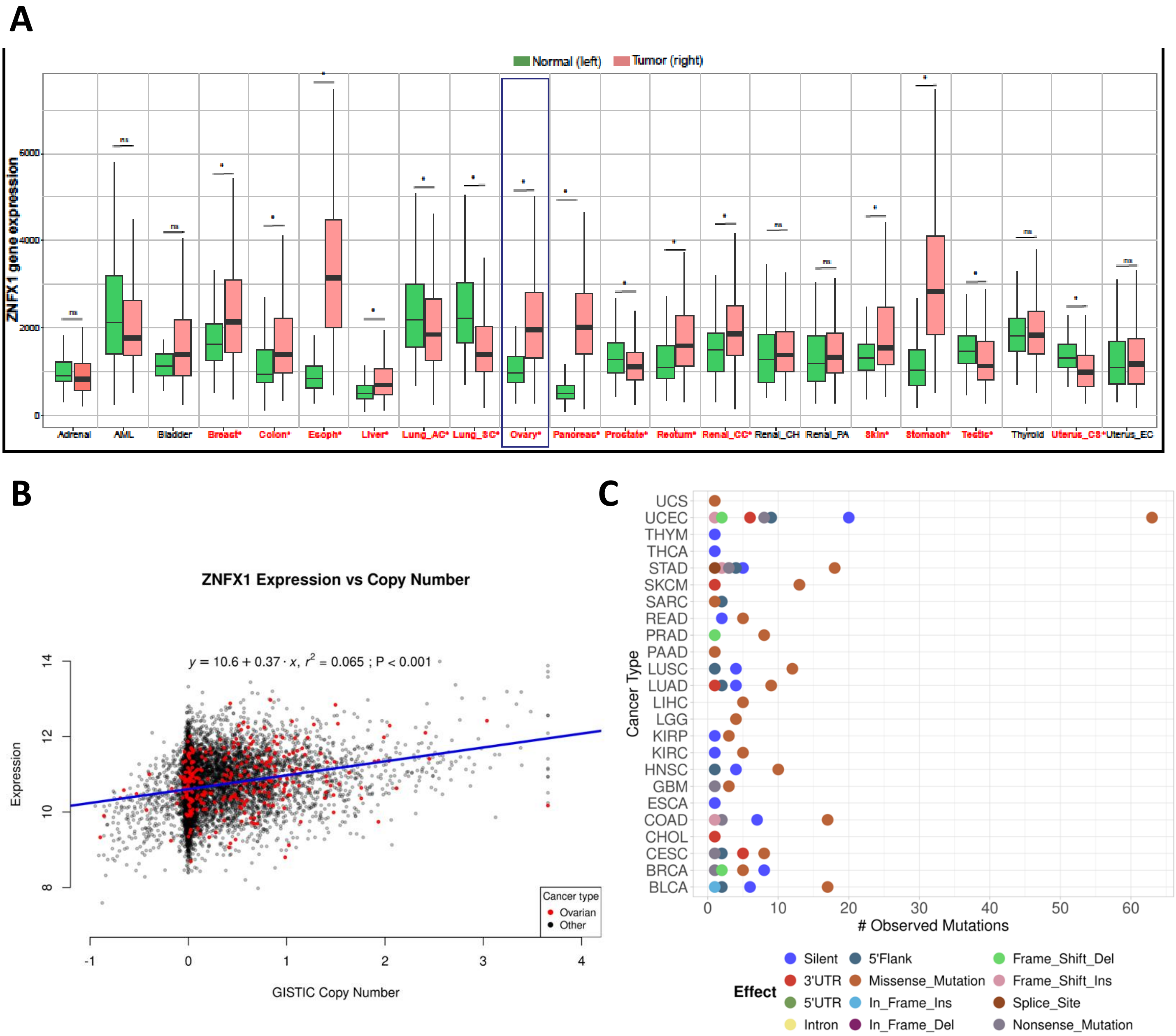


**Supplemental Figure S1 A,** The boxplot shows an expression of ZNFX1 in normal and tumor bulk RNA-seq dataset derived from NCBI GEO, GTex, TCGA, and TARGET databases. In the box-and-whisker plots, the horizontal lines mark the median, the box limits indicate the 25th and 75th percentiles, and the whiskers extend to 1.5× the interquartile range from the 25th and 75th percentiles. The statistical testing of expression levels of signature genes between the groups was performed using two-tailed unpaired Wilcoxon test. **B,** ZNFX1 expression shows a positive relationship to ZNFX1 copy number in pan-cancer tumor samples. Log2 normalized expression and GISTIC gene-level copy number data was obtained from UCSC Xena browser for all TCGA Pan-Cancer solid tumor samples. A linear model was fit on all samples to predict ZNFX1 expression as a function of ZNFX1 copy number. The model explains a statistically significant and weak proportion of variance ($R2 = 0.06$, $F(1, 9438) = 652.08$, $p < .001$, adj. $R2 = 0.06$). The effect of copy number is statistically significant and positive (beta = 0.37, 95% CI [0.34, 0.40], $t(9438) = 25.54$, $p < .001$; Std. beta = 0.25, 95% CI [0.23, 0.27]). Ovarian cancer samples are highlighted in red. Samples from all other cancer types shown in black. **C,** Overall number of observed mutations in ZNFX1 by predicted effect across TCGA cancer types. TCGA Pan-cancer mutation calls were obtained from UCSC Xena browser. The total number of mutations across all samples of each predicted effect type were summed within each cancer type and are shown on the x-axis colored by the predicted effect.

### Supplemental Figure S2

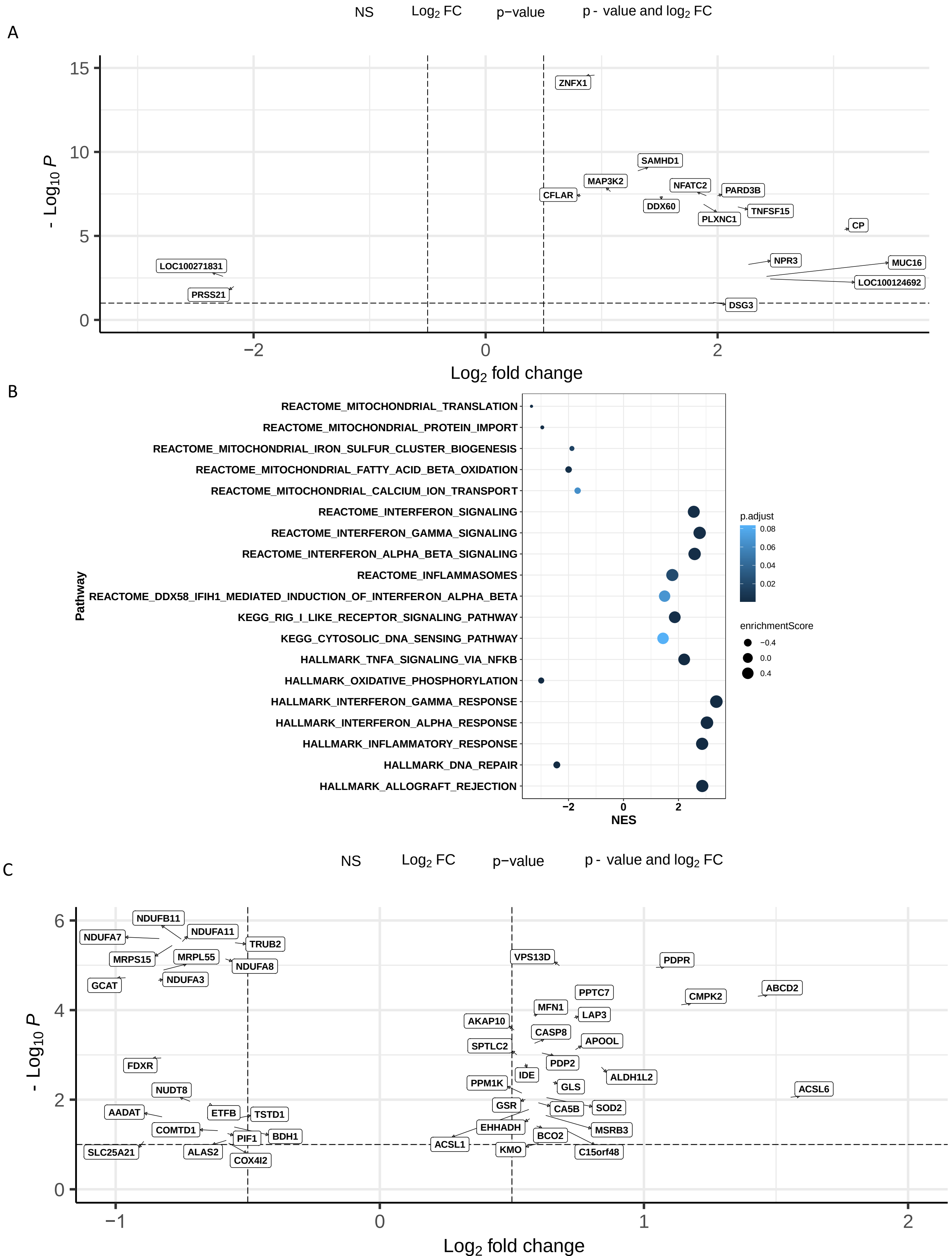

### Supplemental Figure S2

**Supplemental Figure S2**. **A,** Volcano plot for RNAseq differential expression analysis of TCGA triple-negative breast cancer, annotated HGNC symbols, x-axis: log2 fold change in expression: ZNFX1 above median vs. ZNFX1 below median, y-axis: -log10 of FDR controlled adjusted p-value (padj), color mapping: gray: padj> 0.10 and log2 fold change < |0.5|, black: padj< 0.10 and log2 fold change < |0.5|, blue: padj> 0.10 and log2 fold change > |0.5|, and orange: padj< 0.10 and log2 fold change > |0.5|. **B,** Pathway dot plot depicting result gene set enrichment analysis TCGA triple-negative breast cancer on pre-ranked gene list derived from ZNFX1 above median vs. ZNFX1 below median differential expression analysis. Pathways depicted are derived from manual curation of Interferon, Mitochondria, and DNA repair pathways compiled from MSigDB: HALLMARK, KEGG, and REACTOME. x-axis: normalized enrichment score, dot size: enrichment score, color gradation: FDR controlled adjusted p-value. **C,** Volcano plot for RNAseq differential expression analysis TCGA triple-negative breast cancer, MITOCARTA 3.0 symbols, x-axis: log2 fold change in expression: ZNFX1 above median vs. ZNFX1 below median, y-axis: -log10 of FDR controlled adjusted p-value (padj), color mapping: gray: padj> 0.10 and log2 fold change < |0.5|, black: padj< 0.10 and log2 fold change < |0.5|, blue: padj> 0.10 and log2 fold change > |0.5|, and orange: padj< 0.10 and log2 fold change > |0.5|.

### Supplemental Figure S3

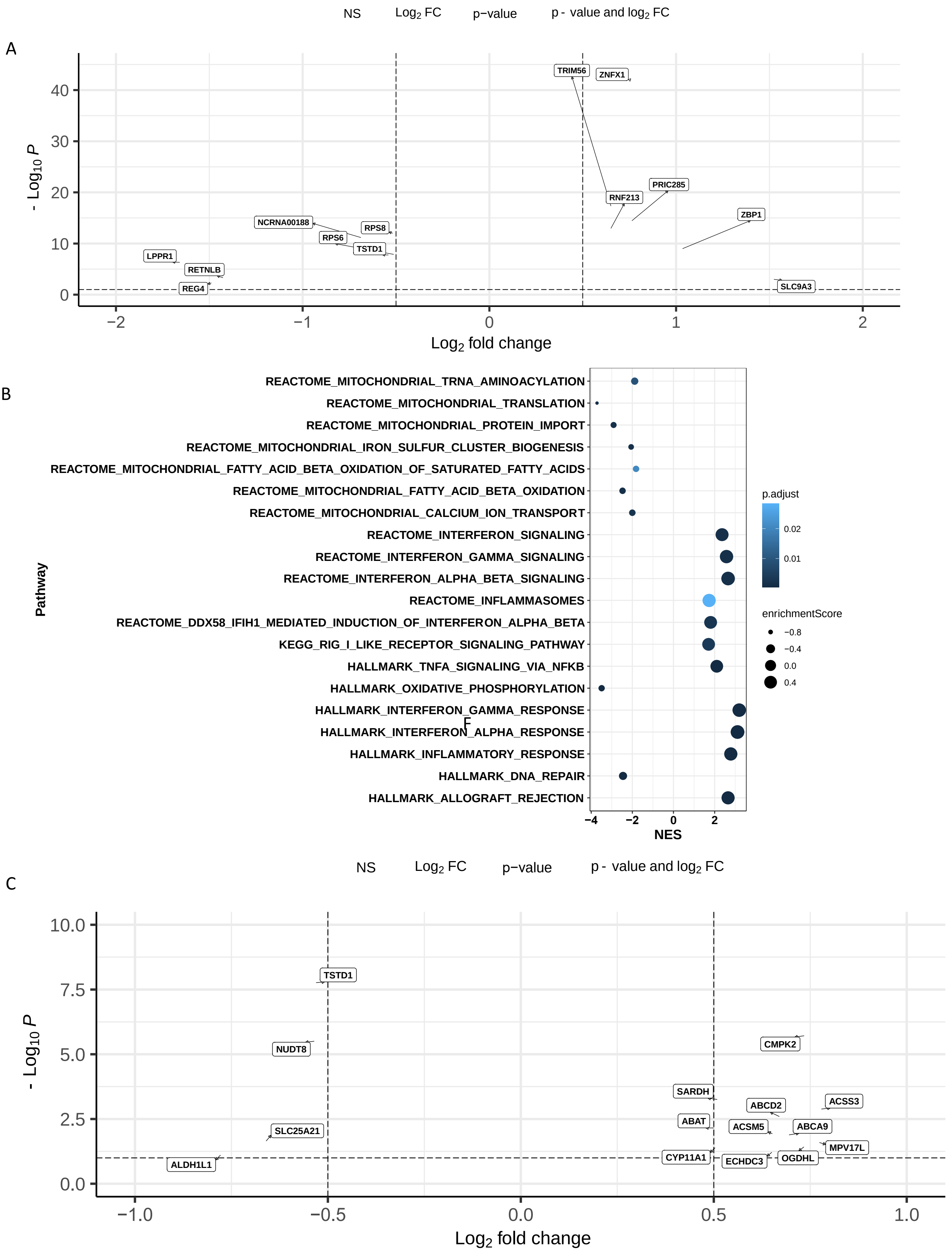

### Supplemental Figure S3

**Supplemental Figure S3**. **A,** Volcano plot for RNAseq differential expression analysis of TCGA colon adenocarcinoma, annotated HGNC symbols, x-axis: log2 fold change in expression: ZNFX1 above median vs. ZNFX1 below median, y-axis: -log10 of FDR controlled adjusted p-value (padj), color mapping: gray: padj> 0.10 and log2 fold change < |0.5|, black: padj< 0.10 and log2 fold change < |0.5|, blue: padj> 0.10 and log2 fold change > |0.5|, and orange: padj< 0.10 and log2 fold change > |0.5|. **B,** Pathway dot plot depicting result gene set enrichment analysis TCGA colon adenocarcinoma on pre-ranked gene list derived from ZNFX1 above median vs. ZNFX1 below median differential expression analysis. Pathways depicted are derived from manual curation of Interferon, Mitochondria, and DNA repair pathways compiled from MSigDB: HALLMARK, KEGG, and REACTOME. x-axis: normalized enrichment score, dot size: enrichment score, color gradation: FDR controlled adjusted p-value. **C,** Volcano plot for RNAseq differential expression analysis TCGA colon adenocarcinoma , MITOCARTA 3.0 symbols, x-axis: log2 fold change in expression: ZNFX1 above median vs. ZNFX1 below median, y-axis: -log10 of FDR controlled adjusted p-value (padj), color mapping: gray: padj> 0.10 and log2 fold change < |0.5|, black: padj< 0.10 and log2 fold change < |0.5|, blue: padj> 0.10 and log2 fold change > |0.5|, and orange: padj< 0.10 and log2 fold change > |0.5|.

### Supplemental Figure S4

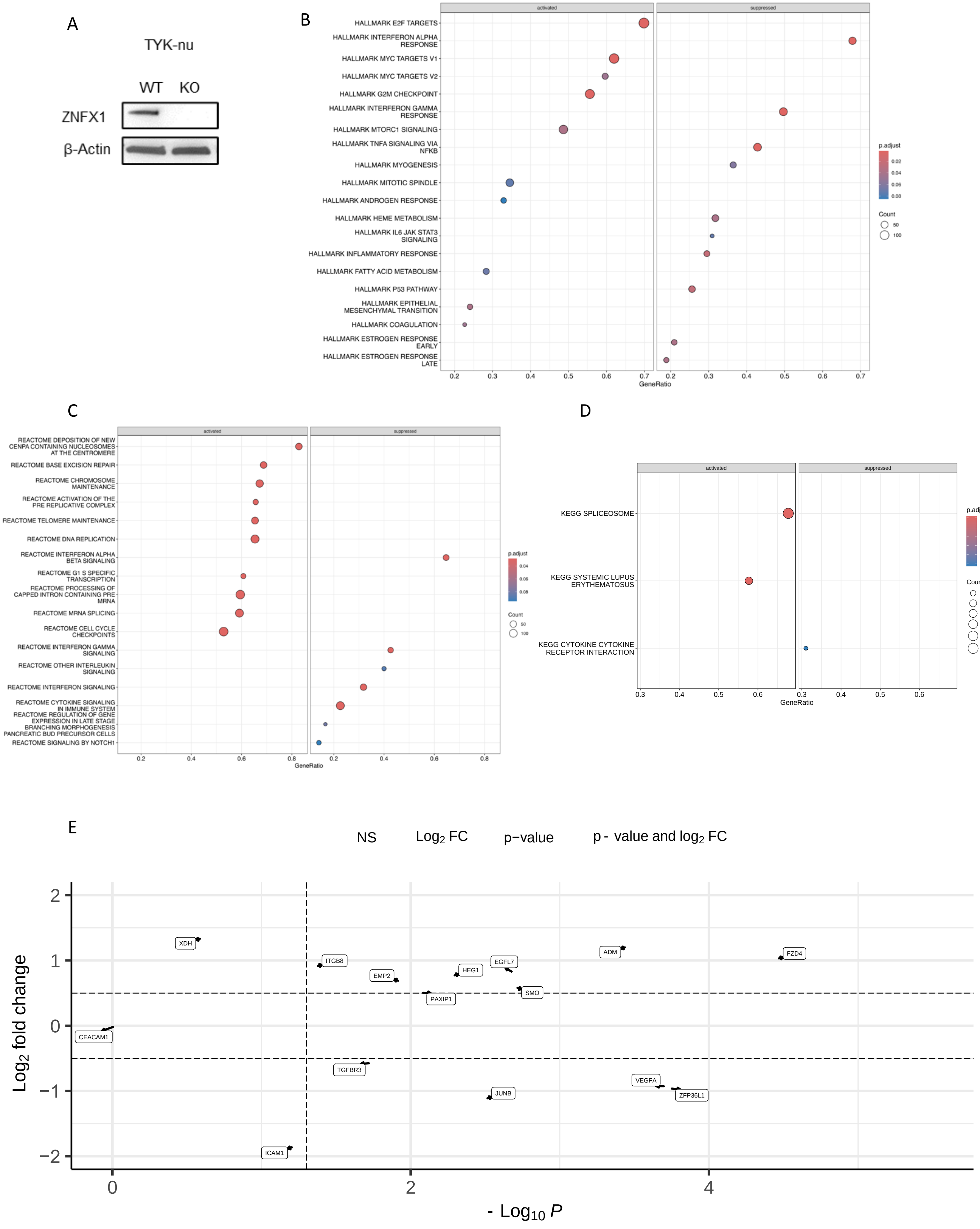

### Supplemental Figure S4

**Supplemental Figure S4: A,** Immunoblotting for ZNFX1 in WT and single clones of ZNFX1 CRISPR KO TYK-nu cells. **B,C,D,** Pathway dot plot depicting top pathway results derived from gene set enrichment analysis on ZNFX1 KO vs. WT TYK-nu cells, (B) HALLMARK pathways, (C) REACTOME pathways, (D) KEGG pathways. x-axis: GeneRatio, dot size: count, color gradation: FDR controlled adjusted p-value. **E,** Volcano plot for RNAseq differential expression analysis of curated vasculogenesis genes for: ZNFX1 KO vs. ZNFX1 WT, x-axis: -log10 of FDR controlled adjusted p-value (padj), y-axis: log2 fold change in expression, color mapping: gray: padj> 0.10 and log2 fold change < |0.5|, black: padj< 0.10 and log2 fold change < |0.5|, blue: padj> 0.10 and log2 fold change > |0.5|, and orange: padj< 0.10 and log2 fold change > |0.5|.

### Supplemental Figure S5

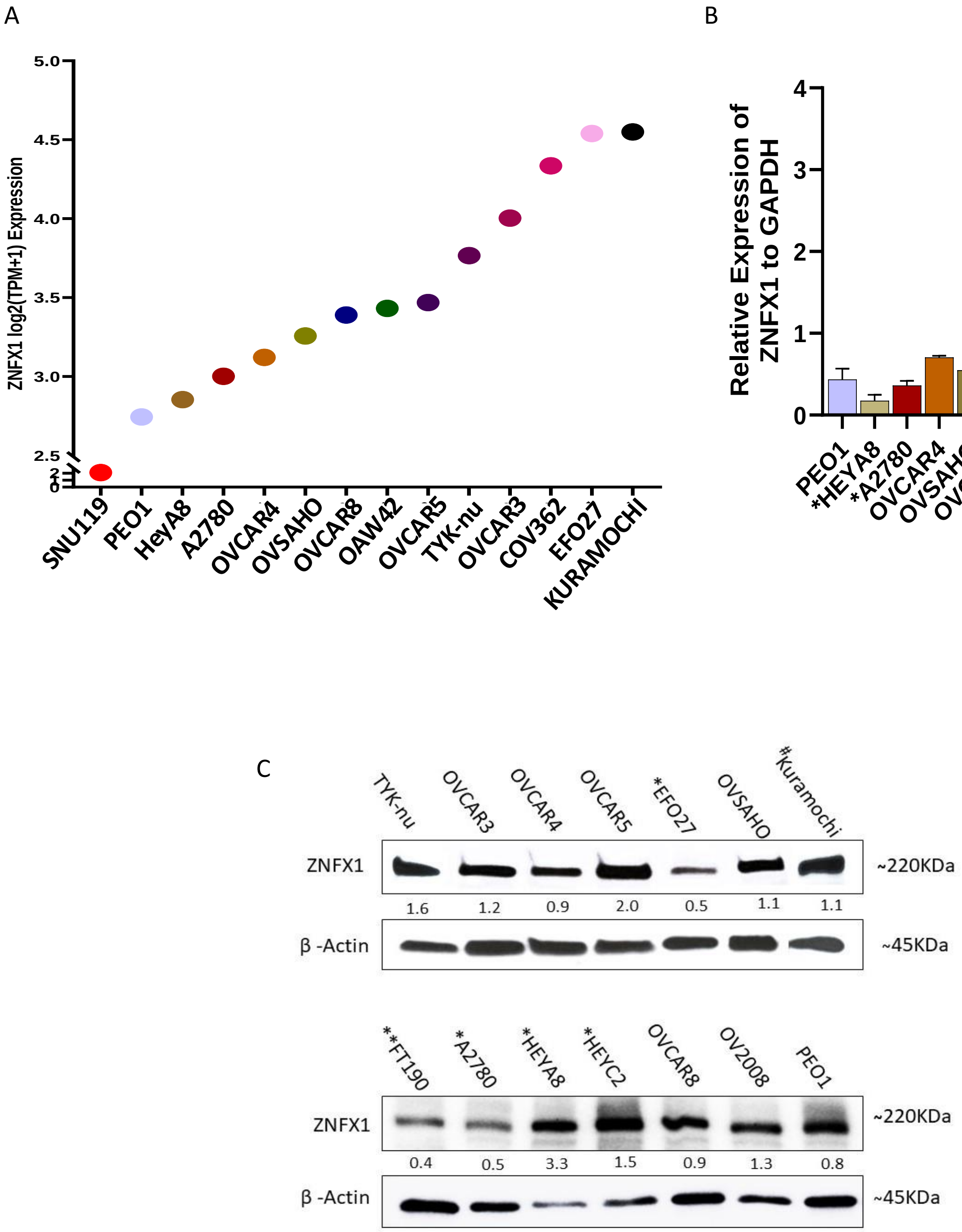


**Supplemental Figure S5:** Relative ZNFX1 gene expression **A,** CCLE database**, B,** RT-qPCR and **C,** Protein expression (western blot analysis) of high grade serous and non-serous (*) ovarian cancer cell lines and normal fallopian tube epithelial cells (**). For all cell lines, 10 µg protein used except for Kuramochi (#; 20 µg of protein was used).

### Supplemental Figure S6

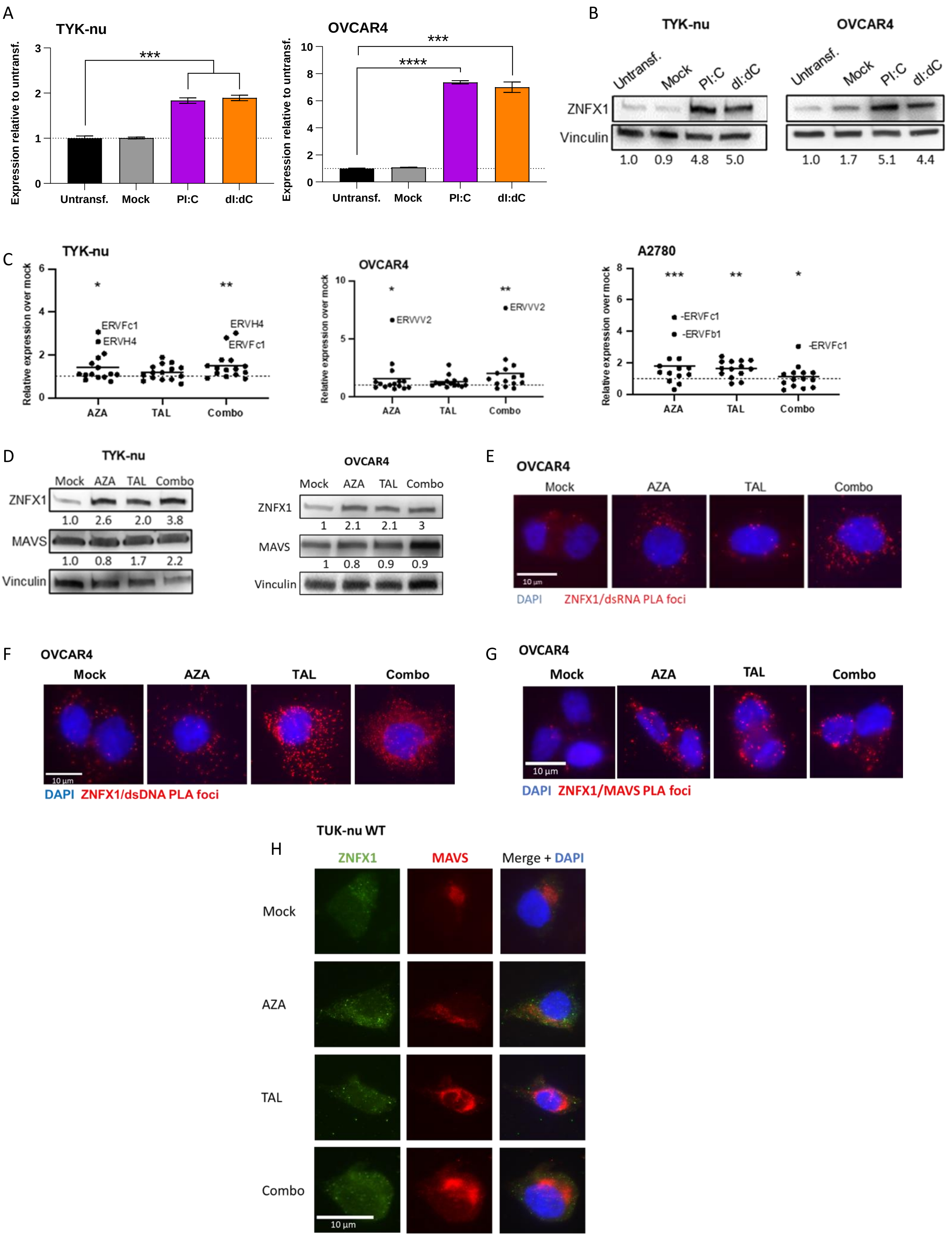

### Supplemental Figure S6

**Supplemental Figure S6. A,B,** Relative expression of ZNFX1 transcripts by qPCR (A) and proteins (B) in TYK-nu and OVCAR4 cells 72 hours after transfection with 1µg poly I:C or poly dI:dC**. C,** Relative expression of ERV gene transcripts by qPCR in TYK-nu, OVCAR4 and A2780 cells after treatment with AZA, TAL, or combination therapy for 6 days. **D,** ZNFX1 and MAVs protein expression levels by immunoblotting following 6 days of AZA (100nM and 150nM), TAL (2.5 and 10nM) and combination treatment in TYK-nu and OVCAR4 cells. **E-G,** Representative immunofluorescence images of ZNFX1 colocalization with dsRNA (E), dsDNA (F), and MAVS (G) by proximity ligation assays in OVCAR4 cells after mock, AZA, TAL and combination treatment. **H,** Representative immunofluorescence images showing ZNFX1 and MAVS colocalization in TYK-nu following 6 days treatment with AZA, TAL, or combination. Data are presented as mean ± SEM with p values derived from two-tailed unpaired Student's t-test or ANOVA as appropriate. * $p<0.05$, ** $p<0.01$, *** $p<0.001$.

### Supplemental Figure S7

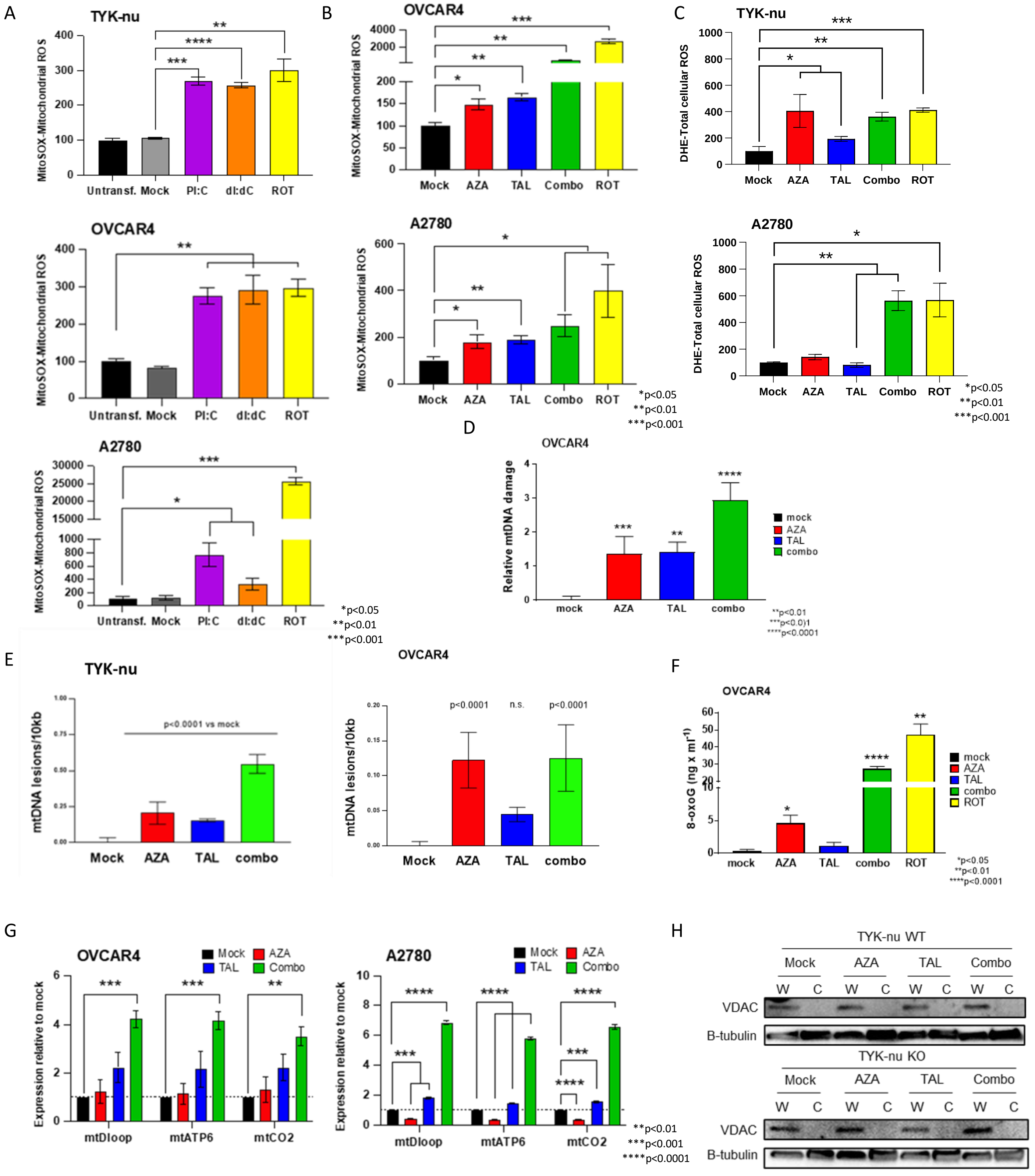

### Supplemental Figure S7

**Supplemental Figure S7.** Increased ZNFX1 leads to increased mt ROS, DNA damage and dsDNA leakage into the cytosol in TYK-nu, OVCAR4 and A2780 ovarian cancer cell lines. **A, B**, mtROS levels measured by flow cytometry of mitosox in treated with 1µg poly I:C or 1µg poly dI:dC 72 hours after transfection (A); After 6 days of treatment with AZA, TAL, and combination therapy (B). **C,** Total cellular ROS levels measured by flow cytometry of DHE in TYK-nu, OVCAR4 and A2780 cells after 6 days of treatment with AZA, TAL and combination therapy in TYK-nu and A2780. **D,** Relative DNA damage as measured by adapted qPCR long PCR method in mtDNA in TYK-nu and OVCAR4 cells treated with AZA, Tal, and combination therapy. **E,** Relative mitochondrial DNA damage detected by end-point long-range PCR in TYK-nu and OVCAR4 following 6 days' treatment with AZA, TAL, or combination (TYK-nu: AZA 100nM, TAL 2.5nM, or combination; OVCAR4: AZA 150nM, TAL 10nM, or combination). **F,** 8OXOG as measured by ELISA in mtDNA from OVCAR4 cells treated with AZA, Tal, and combination therapy. Rotenone used as a positive control. **G,** Relative expression of mtDNA (ATP6/8, mtCO2, MtND1) in the cytosol of OVCAR4 and A2780 cells, following AZA, Tal, and combination therapy, as measured by qPCR. **H,** Western blot showing VDAC and β-tubulin protein expression in whole cell lysate (W) and cytoplasmic fraction (C) in TYK-nu and TYK-nu KO following 6 days' treatment with AZA, TAL, or combination All data are presented as mean ± SEM with statistical significance p values derived from two-tailed unpaired Student's t test (or ANOVA). Rotenone is used as a positive control.

### Supplemental Figure S8

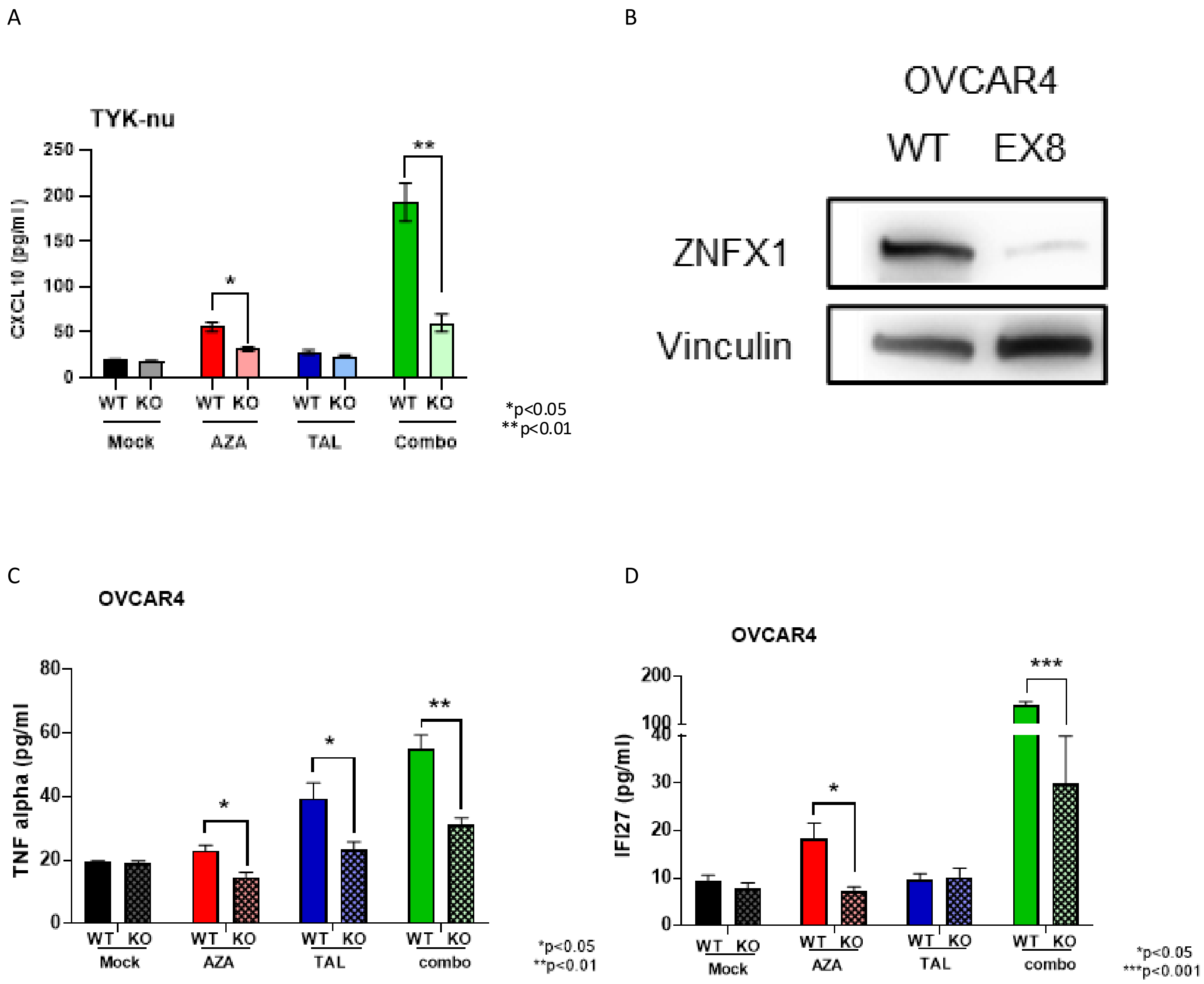


**Supplemental Figure S8. A,** Levels of cytokine CXCL10 measured by ELISA assays after mock, AZA, 2.5 nM Tal, or AZA/Tal combination treatment in TYK-nu (parental and ZNFX1 KO) cells. **B**, Immunoblotting for ZNFX1 in bulk population of OVCAR4 cells following nucleofection with CRISPR gRNAs targeting exon 8 of ZNFX1. **C, D,** Levels of cytokines, TNFα (B) and IFI27 (C) measured by ELISA assays after mock, AZA, 2.5 nM Tal, or AZA/Tal combination treatment in OVCAR4 (parental and ZNFX1 KO) cells in bulk population of OVCAR4 cells. All data are presented as mean ± SEM with statistical significance p values derived from two-tailed unpaired Student's t test (or ANOVA).

#### Supplemental Figure S9

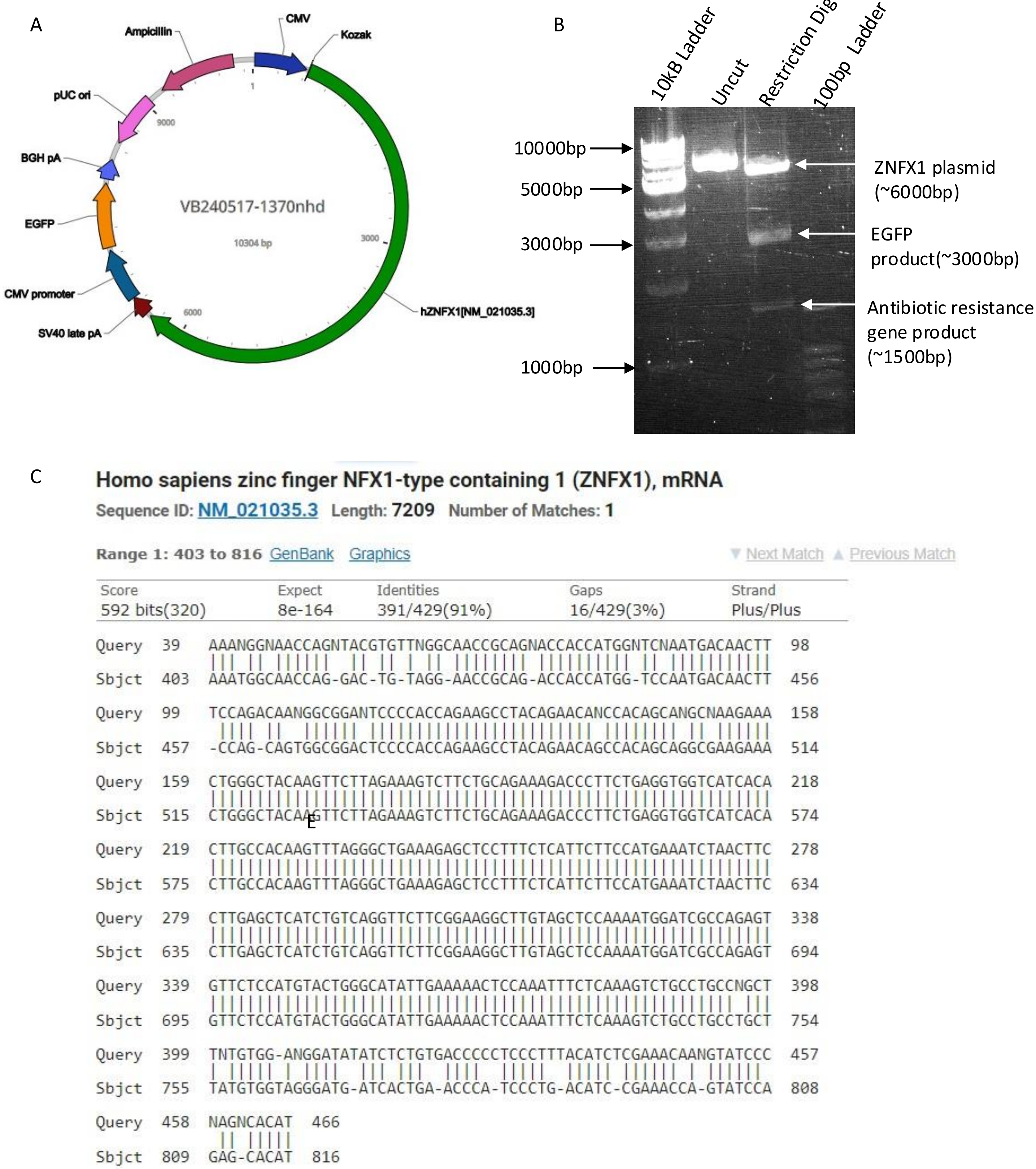


**Supplementary Figure S9**. ZNFX1 rescue experiments. **A,** Gene construct. Vector map showing ZNFX1 insert locations along with reporter and antibiotic resistance gene locations. **B**, Agarose gel electrophoresis of plasmid DNA digested with BsgR1 restriction enzyme. The digestion resulted in three distinct fragments: the largest fragment (~6000 bp), corresponding to the ZNFX1 gene insert, and two smaller fragments. **C**, The nucleotide blast shows that ZNFX1 cDNA sequence in plasmid align with human ZNFX1

### Supplemental Figure S10

A

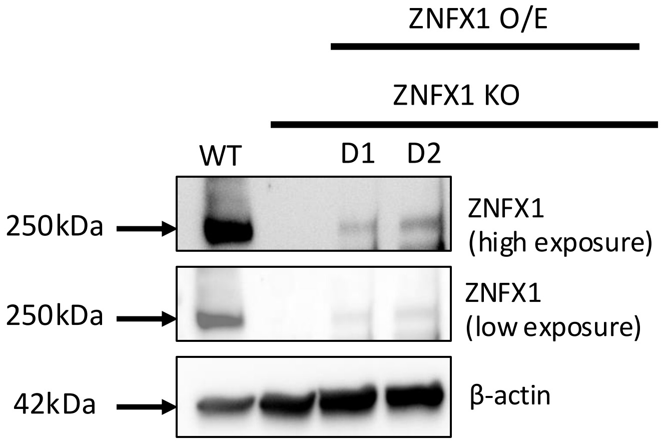


B

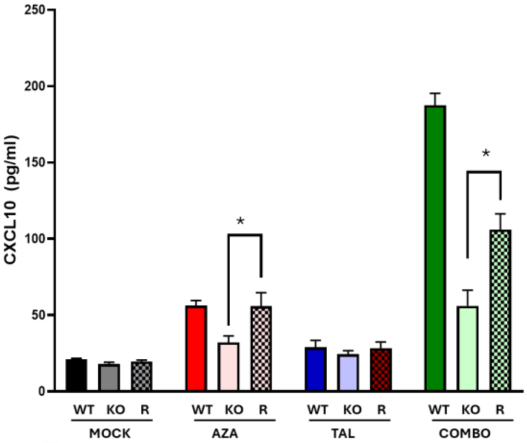


**Supplementary Figure S10**. ZNFX1 rescue experiments. **A,** Western blot analysis following transient transfection of WT ZNFX1 plasmid in ZNFX1 KO cells following 2 days (D1, D2). **B**, Levels of cytokine CXCL10 in rescue experiment overexpressing ZNFX1 plasmid construct (R), with AZA, 2.5 nM Tal, or AZA/Tal combination treatment in ZNFX1 KO TYK-nu cells, measured by ELISA assays. All data are presented as mean ± SEM with p values derived from two-tailed unpaired Student's t-test or ANOVA as appropriate. * p<0.05, ** p<0.01, *** p<0.001.

### Supplemental Figure S11

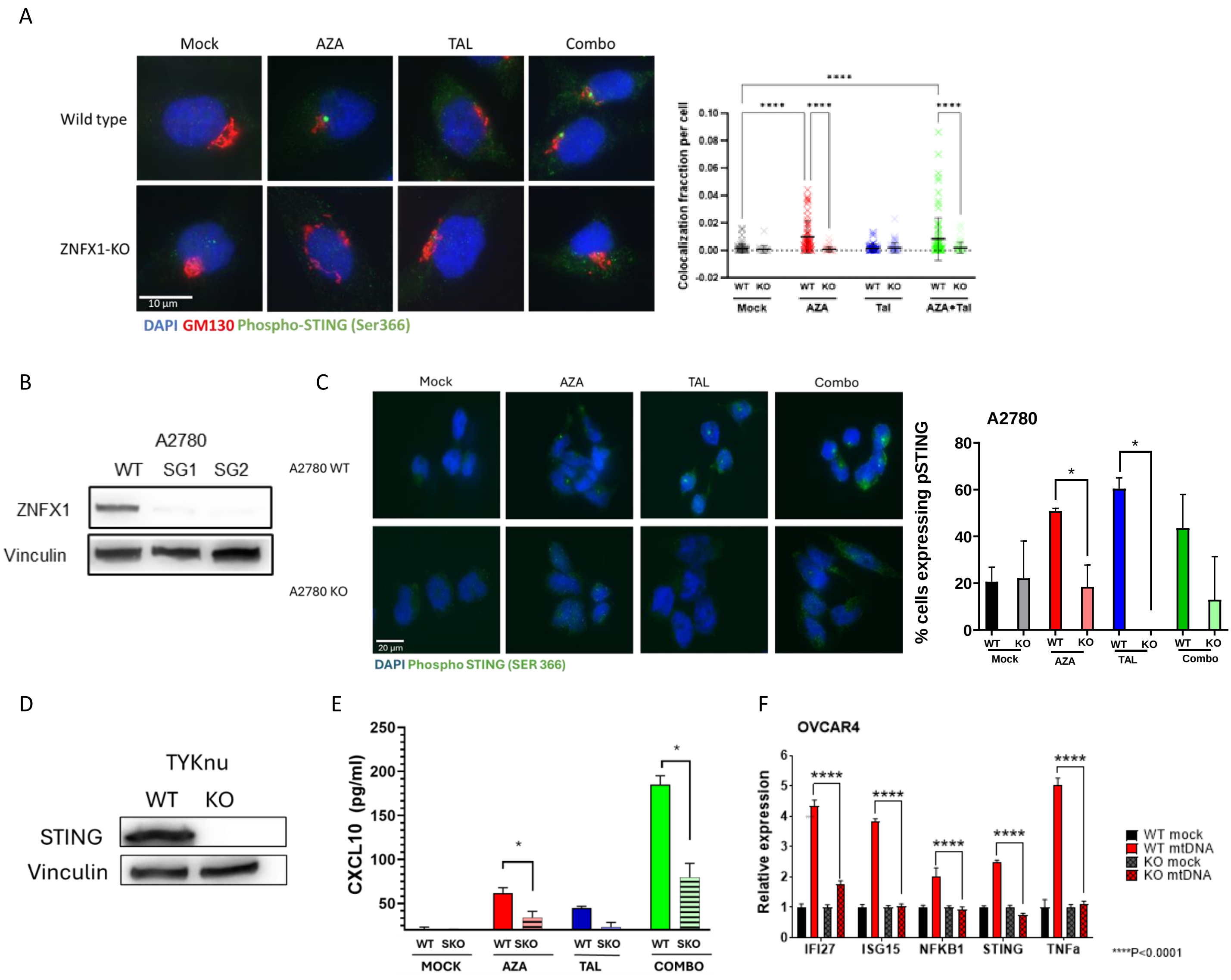


**Supplemental Figure S11.** STING activity and KO experiments. **A,** Left, representative immunofluorescence images of phosphoSTING (green fluorescence) and Golgi protein GM130 (red fluorescence) in parental and ZNFX1 KO OVCAR4 cells following treatment with AZA, TAL, or combination. Right, Graph quantitating images. **B,** Immunoblotting for ZNFX1 in parental and KO A2780 cells. Vinculin used as a loading control. C, representative immunofluorescence images of phosphoSTING (green fluorescence) in A2780 parental and ZNFX1 KO cells following treatment with AZA, TAL, or combination. Right, Graph quantitating images. **D,** Immunoblotting for ZNFX1 in parental and STING KO TYK-nu cells. Vinculin used as a loading control. **E,** Levels of cytokine CXCL10 in TYK-nu STING KO (SKO) and TYK-nu WT cell lines with AZA, 2.5 nM Tal, or AZA/Tal combination treatment., measured by ELISA assays. **F,** Relative RNA expression of IFN/inflammasome genes (IFI27, ISG15, NFKB1, STING, TNFa) by qPCR 72hrs after transfection of mt DNA in OVCAR4 (parental and ZNFX1 KO) cells. All data are presented as mean ± SEM with statistical significance p values derived from two-tailed unpaired Student’s t test (or ANOVA).

### Supplemental Figure  S12

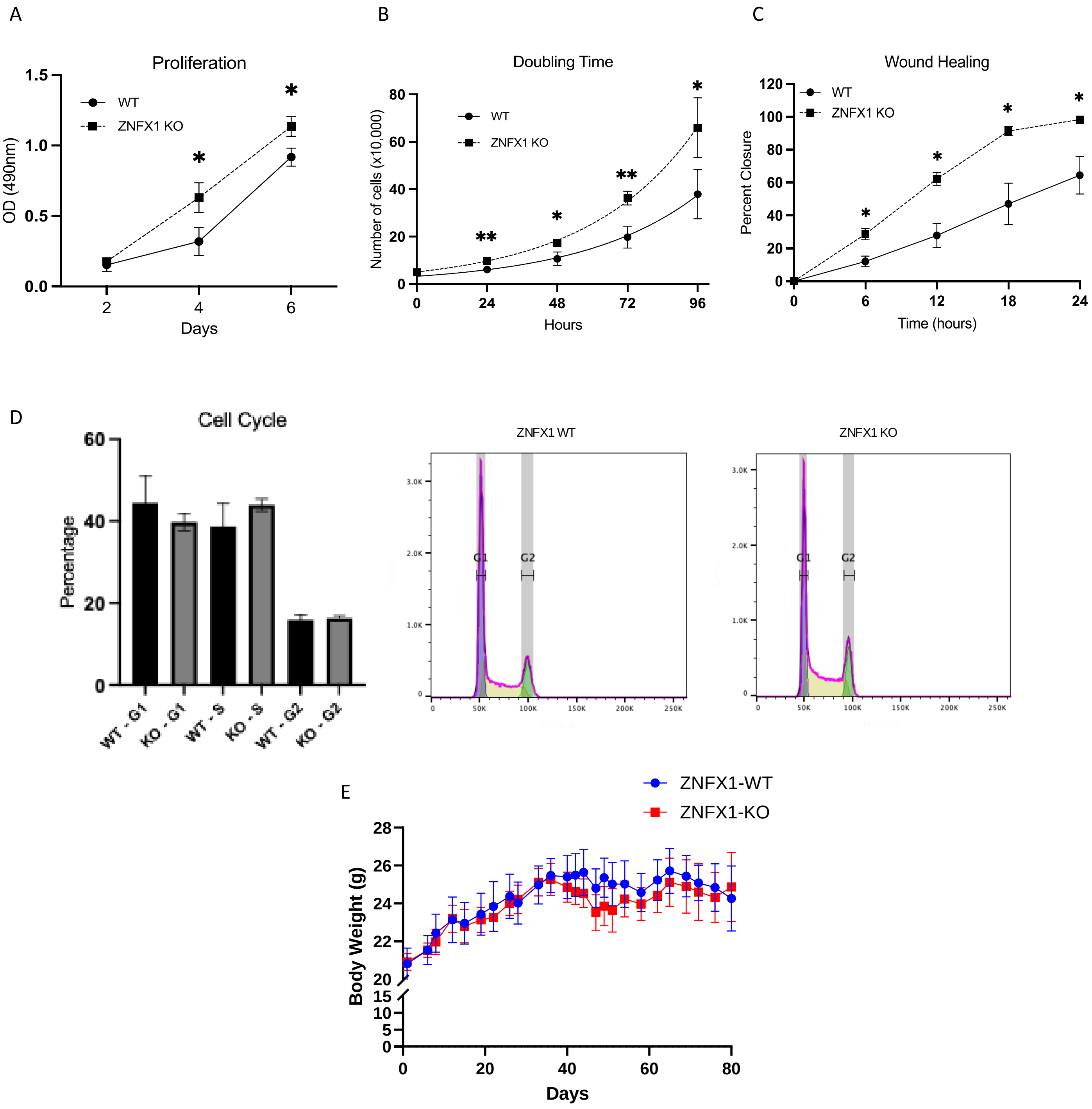


**Supplemental Figure S12**. Functional analysis of TYK-nu WT and ZNFX1 KO cells. **A,** Proliferation (seeding density 1000 cells), **B,** Cell doubling time and **C,** Wound closure (percentage over 24 hours). **D,** Cell cycle (propidium iodide labeling and flow cytometry analysis). Left, representative population histograms of the cell-cycle phase distribution for WT and ZNFX1-KO; blue, G0/G1 phase; yellow, S phase; green.  Right, G2/M phase. **E,** Body weights (grams) of NSG mice growing subcutaneous xenografts of TYK-nu ZNFX1-WT  or -KO cells. All data are presented as mean ± SEM with p values derived from two-tailed unpaired Student's t test or ANOVA as appropriate. * p<0.05, ** p<0.01, *** p<0.001, **** p<0.0001.

### Supplemental Figure S13

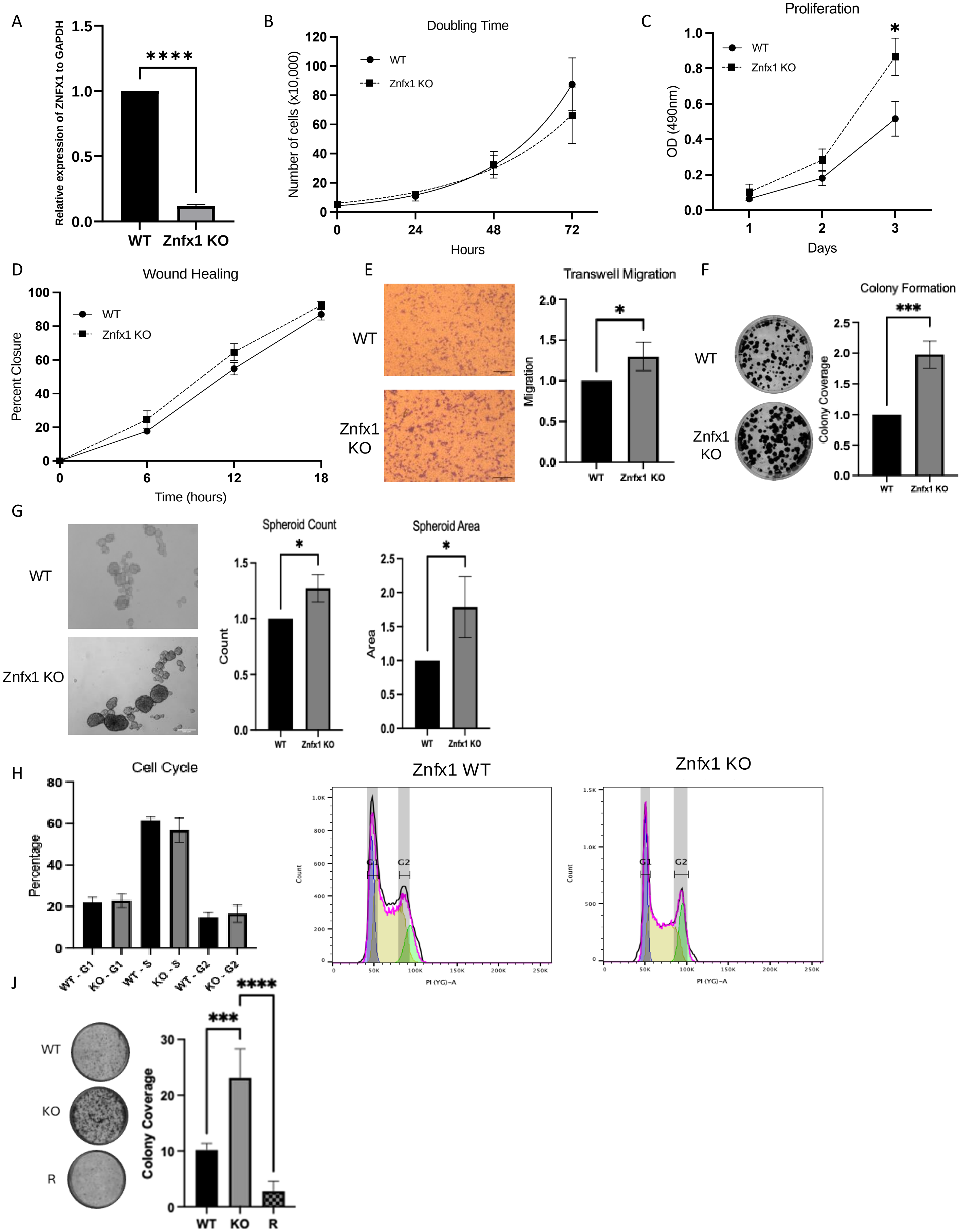

**Supplemental Figure S13**. Functional analysis KPCA-WT or ZNFX1-KO cells (**A)** Relative expression (RT-qPCR), **B,** Cell doubling time, **C,** Proliferation, **D,** Wound healing, **E,** Cell migration (representative images (left), graphical representation (right), **F,** Clonogenic survival assays (500 cells per well; representative Images of colonies (left) and graphs of colony coverage (right), **G,** Spheroid formation assays (3000 cells/well; representative images are shown (left)). Spheroid formation was assessed through microscopy. Quantification was performed using Image J, with data presented as both spheroid area and spheroid count (middle and right). **H,** Cell cycle (propidium iodide labeling and flow cytometry analysis; *left,* representative population histograms of the cell-cycle phase distribution, blue, G0/G1 phase; yellow, S phase; green; *right,* G2/M phase), **J**, Clonogenic survival assays with overexpression of WT ZNFX1 in ZNFX1 KO cells (1000 cells per well; representative images of colonies (left) and graphs of colony coverage (right)). All data are presented as mean ± SEM with p values derived from two-tailed unpaired Student's t test or ANOVA as appropriate. * $p<0.05$, ** $p<0.01$, *** $p<0.001$, **** $p<0.0001$

### Supplemental Figure S14

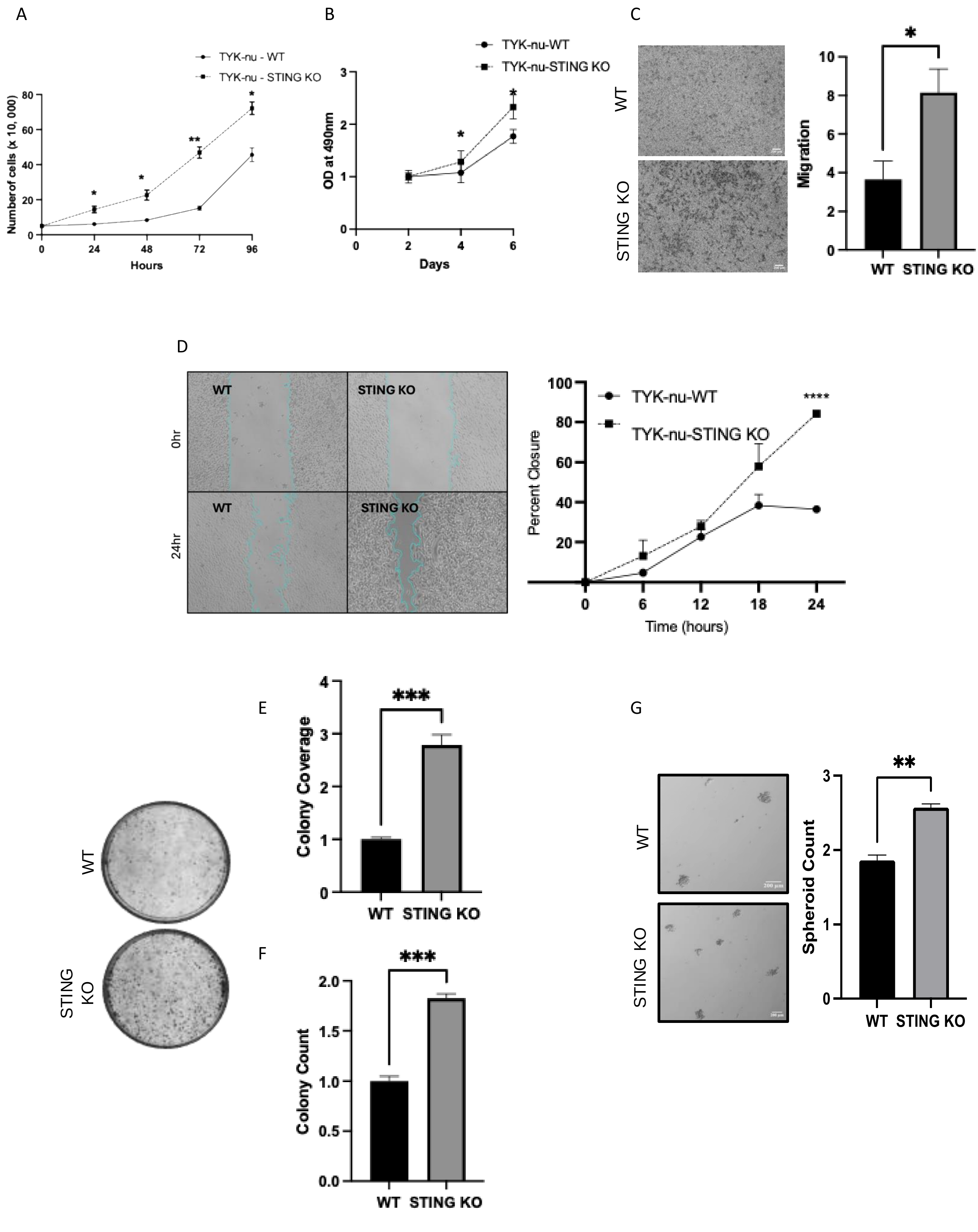

**Supplemental Figure S14**. Functional analysis of WT and STING KO TYK-nu cells **A,** Cell doubling time, **B,** Proliferation, **C,** Cell migration (representative images (left), graphical representation (right), **D**, Wound healing, **E, F,** Clonogenic survival assays (1000 cells per well; representative Images of colonies (left) and graphs of colony coverage (right), **G,** Spheroid formation assays (3000 cells/well; representative images are shown (left)). Spheroid formation was assessed through microscopy. Quantification was performed using Image J, with data presented as both spheroid area and spheroid count (middle and right). All data are presented as mean ± SEM with p values derived from two-tailed unpaired Student's t test or ANOVA as appropriate. * p<0.05, ** p<0.01, *** p<0.001, **** p<0.0001

### Supplementary Figure S15

A

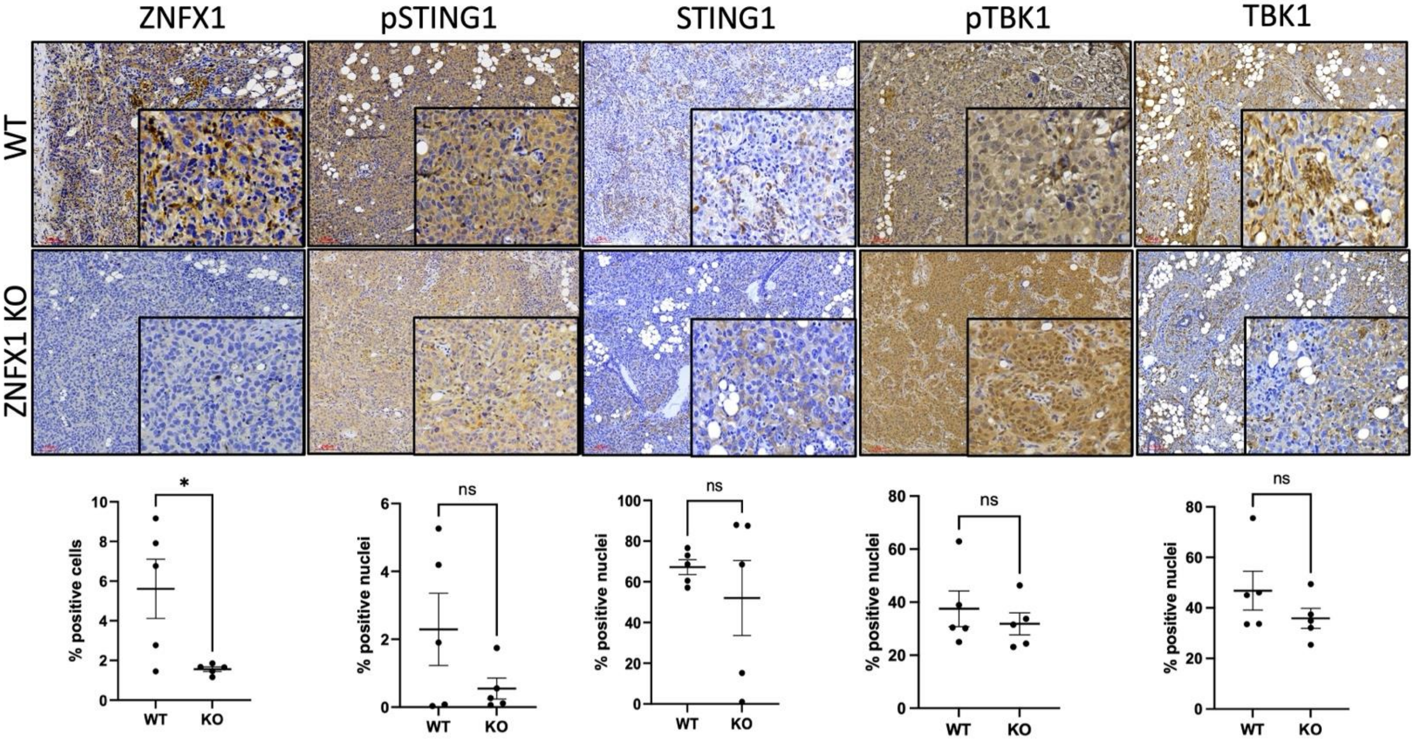


B

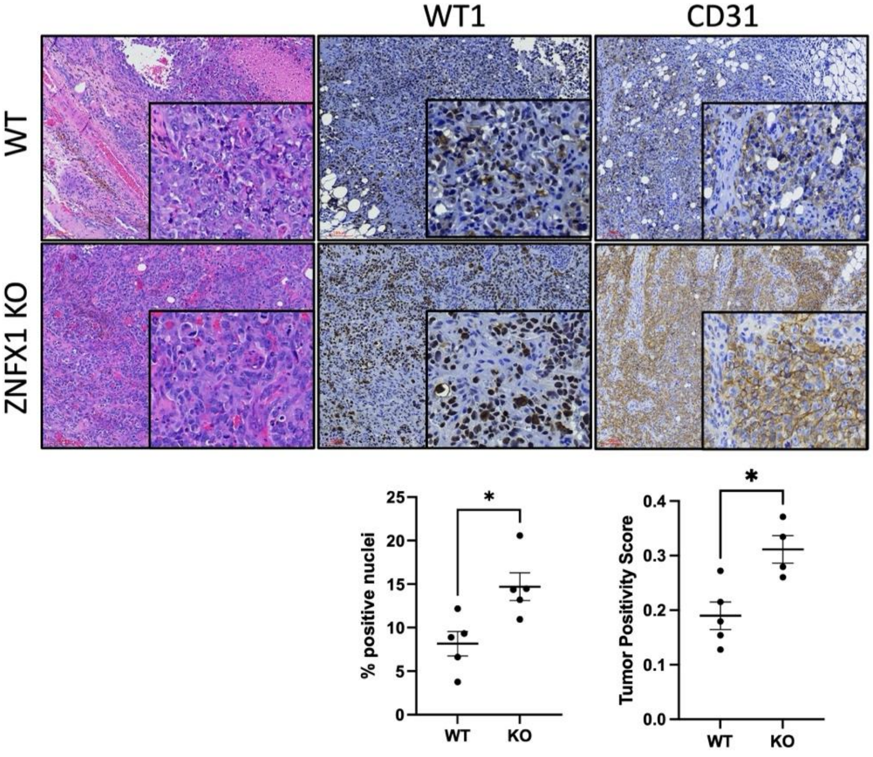


**Supplementary Figure S15**. Representative images of hematoxylin and eosin staining of xenograft tumors and with specific antibodies against **A,** ZNFX1, pSTING, STING, pTBK1, TBK1, and **B,** WT1 and CD31. The respective quantification is given below. Images are shown at 40x. Bar size, 100 μm. All data are presented as mean ± SEM with p values derived from two-tailed unpaired Student's t-test or ANOVA as appropriate. * p<0.05, ** p<0.01, *** p<0.001.

#### Supplemental Figure S16

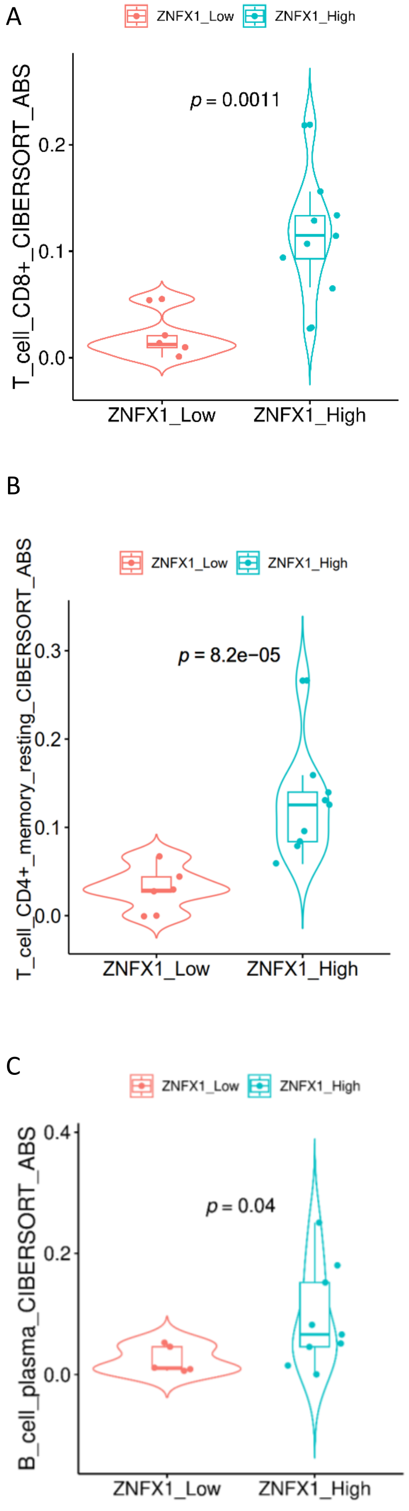


.

**Supplementary Figure S16.** Violin plots showing correlations between ZNFX1 expression (X axis) and predicted absolute proportions (Y axis) of T cells (**A,** CD8+; **B,** CD4+) and B cells (**C,** plasma). Statistical significance was assessed using Wilcoxon rank-sum tests between the ZNFX1 groups.

#### Supplemental Figure S17

A

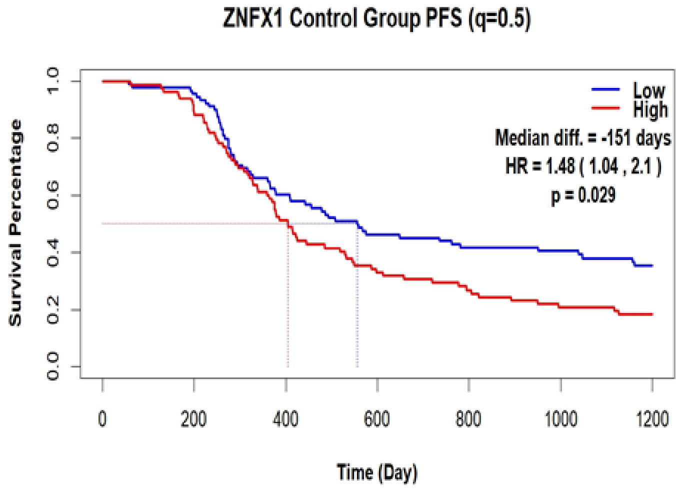


B

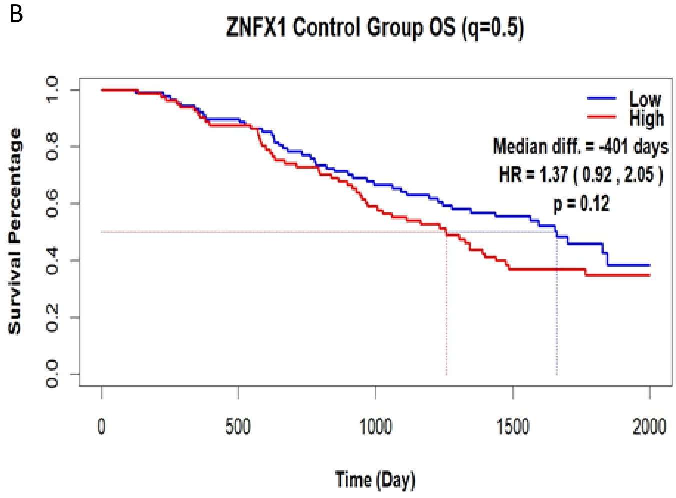


**Supplemental Figure S17.** ICON7 trial data KM plots of **A,** PFS and **B**, OS rates of low and high ZNFX1 expression among patients in the ICON7 trial receiving standard chemotherapy treatment (N=181). High and low expression are determined by median expression level. HR = hazard ratio (estimate (95% CI)). Lower expression of ZNFX1 is consistent with improved PFS, and OS may also be improved.

#### Supplemental Figure S18

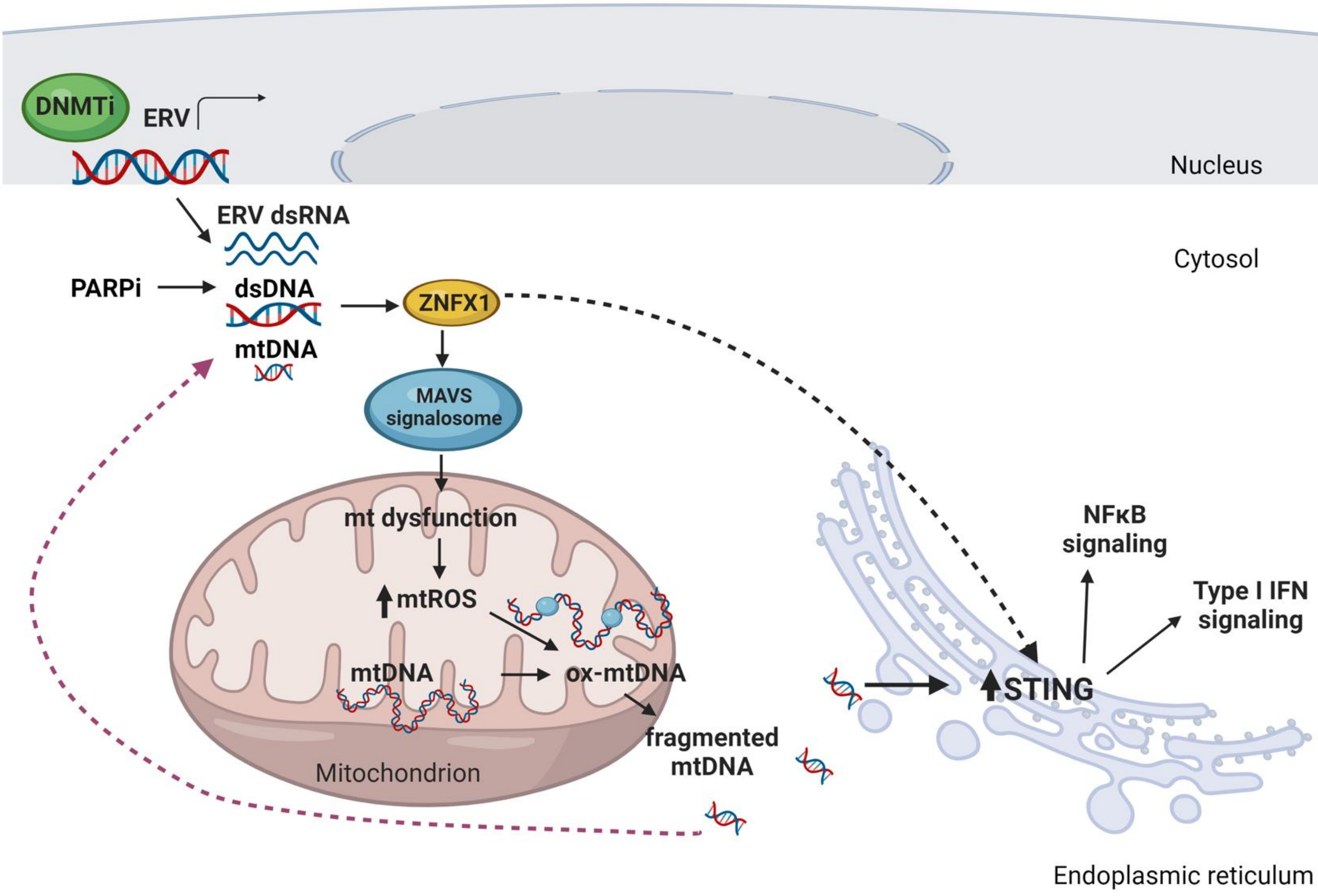


**Supplemental Figure S18.**

Model for ZNFX1 induction of mt dysfunction and inflammatory signaling triggered by DNMTi and PARPi therapy in OC. DNMTis increase expression of ERV transcripts which accumulate as cytosolic dsRNA. PARPi treatment produces DNA damage and fragmentation that accumulates as cytosolic dsDNA. ZNFX1 senses dsRNA/DNA, localizes to MAVs on mitochondrial membrane, and produces mt dysfunction that increases mtROS, oxidative mtDNA damage, mtDNA fragmentation, and mtDNA leak into the cytosol. Cytosolic mtDNA activates STING-dependent interferon and inflammasome signaling. ZNFX1 may also play a role in detecting mtDNA in the cytosol (red dashed line), leading to STING activation, as indicated by the dashed arrow. Created with Biorender.com.
