## Supplemental Materials and Methods for "ZNFX1 is a Novel Master Regulator in Epigenetically-induced Pathogen Mimicry and Inflammasome Signaling in Cancer"

**Supplementary Materials and methods:**

Cell lines, culture conditions and reagents:

The KPCA cell lines were developed and described in Iyer et al[1]. Epithelial ovarian cancer cell lines (CP70, A2780, HeyC2, C272) were maintained in RPMI-1640 (Invitrogen, Carlsbad, CA) supplemented with 10% fetal bovine serum (FBS) (ATCC, Manassas, VA, USA) and 1% Penicillin-Streptomycin Solution (ATCC, Manassas, VA, USA). OVCAR8, OV2008, HEYA2 were maintained in DMEM (Invitrogen, Carlsbad, CA) supplemented with 10% fetal bovine serum (FBS) (ATCC, Manassas, VA, USA) and 1% Penicillin-Streptomycin Solution (ATCC, Manassas, VA, USA). Cell lines were tested for mycoplasma contamination (Manassas, VA, USA). To ensure cell line integrity, all cell lines were thawed at frequent intervals and not used beyond 40 passages. Additionally, cell morphology was monitored for each cell line, and proper media and growth conditions selected.

**Table 1.**

| Name | Sequence |
|---|---|
| F-Actin | CACCATTGGCAATGAGCGGTTC |
| R-Actin | AGGTCTTTGCGGATGTCCACGT |
| F-GAPDH | GTCTCCTCTGACTTCAACAGCG |
| R-GAPDH | ACCACCCTGTTGCTGTAGCCAA |
| F-ERV-K1 | ATCCTATGGCACCACCTAGTA |
| R-ERV-K1 | GCCTCAGTATCTCCTTCCTTTC |
| F-ERVV2 | CTTCTTTCTGAGCTCCTGTCTC |
| R-ERVV2 | GTCCTCTGGTTCTTGGTCTTC |
| F-ERVMER34-1 | CCATGGAAGCTCAAGGTCTATC |
| R- ERVMER34-1 | GAAGGGTCCACTGCCATTT |
| F-ERVW1 | CAAGTCCCTTCCCTCTAATTCC |
| R-ERVW1 | TCCACTCCAGCCACTTTAAC |
| F-ERV-FRD 1 | AGCCAGCTCTCAAAGGAAATAG |
| R-ERV-FRD 1 | GAAGGACTACGGCTGCTAAAG |
| F-ERVW2 | CCACTGTCTGTTGGACTTACTT |
| R-ERVW2 | TGGGAGATTGCTTCCTTTACTT |
| F-ERV-H1 | GCCCATTCTCTCTCTCCATATC |
| R-ERV-H1 | CCTGACATTCCTGCCTTCTTA |
| F-ERVFXA34 | CAGGAAACTAACTTTCAGCCAGA |
| R-ERVFXA34 | TAAAGAGGGCATGGAGTAATTGA |
| F-ERV-Fc1 | TACACCCTTACTCCCGTCTT |
| R-ERV-Fc1 | GCCTAACATTCCGACCTCATAC |
| F-ERV-Fc2 | CTGGAAGCTACACACTCCATAC |
| R-ERV-Fc2 | TGCCAAGAGGTGGGTTATTC |
| F-ERV-Fb1 | ATATCCCTCACCACGATCCTAATA |
| R-ERV-Fb1 | CCCTCTGTAGTGCAAAGACTGATA |
| F-ERV-K8 | CCCATCAATCCACCAAGTCTTA |
| R-ERV-K8 | CCTATTTCTTCGGACCTGTTCTT |
| F-ERV-K10 | GTCCCAAGTGTTTCAGGGAATA |
| R-ERV-K10 | GAAGCAGAGAGACTGCTTGTATAG |
| R-IL18 | ACT GGT TCA GCA GCC ATC TT |
| F-IL18 | GGA ATT GTC TCC CAG TGC AT |
| F-MDA5 | GCT GAA GTA GGA GTC AAA GCC C |
| R-MDA5 | CCA CTG TGG TAG CGA TAA GCA G |
| F-RIG1 | CAC CTC AGT TGC TGA TGA AGG C |
| R-RIG1 | GTC AGA AGG AAG CAC TTG CTA CC |
| F-ZNFX1 | GGCAGAGGGAAGAGAGATTTAG |
| R-ZNFX1 | TTCTCCTGGTCATGTCTTTGG |
| F-MAVS | GTC ACT TCC TGC TGA GA |
| R-MAVS | TGC TCT GAA TTC TCT CCT |
| F-CMPK2 | TGGAGACCAGGCATCTTAATTT |
| R-CMPK2 | CTCACTGGAACATGATGAGAGG |
| F-TFAM | GGGAAGGAGGGTTGTGTATTT |
| R-TFAM | AGGAGTTAGCCAAACGCAATA |
| F MT ATP6 | TAGCCCACTTCTTACCACAAGGCA |
| R MT ATP6 | TGAGTAGGTGGCCTGCAGTAATGT |
| F-MT-ND1 | CACCCAAGAACAGGGTTTGT |
| R-MT-ND1 | TGG CCATGGGTATGTTGTTAA |
| F-MT-CO2 | AATCGAGTAGTACTCCCGATTG |
| R-MT-CO2 | TTCTAGGACGATGGGCATGAAA |
| F-MT-D-LOOP | CTATCACCCTATTAACCACTCA |
| R-MT-D-LOOP | TTCGCCTGTAATATTGAACGTA |
